# Blood cultures contain populations of genetically diverse *Candida albicans* strains that may differ in echinocandin tolerance and virulence

**DOI:** 10.1101/2024.10.16.618724

**Authors:** Giuseppe Fleres, Shaoji Cheng, Hassan Badrane, Christopher L. Dupont, Josh L. Espinoza, Darren Abbey, Eileen Driscoll, Anthony Newbrough, Binghua Hao, Akila Mansour, M. Hong Nguyen, Cornelius J. Clancy

**Affiliations:** University of Pittsburgh, Pittsburgh, PA, USA; J. Craig Venter Institute, La Jolla, CA, USA; University of Minnesota, Minneapolis, MN, USA; University of Pittsburgh Medical Center, Pittsburgh, PA, USA; VA Pittsburgh Healthcare System, Pittsburgh, PA, USA

**Keywords:** *Candida albicans*, Bloodstream infections, Genetic diversity, Echinocandin tolerance, Aneuploidy, Chromosome 7 trisomy

## Abstract

It is unknown whether within-patient *Candida albicans* diversity is common during bloodstream infections (BSIs). We determined whole genome sequences of 10 *C. albicans* strains from blood cultures (BCs) in each of 4 patients. BCs in 3 patients contained mixed populations of strains that differed by large-scale genetic variants, including chromosome (Chr) 5 or 7 aneuploidy (*n*=2) and Chr1 loss of heterozygosity (*n*=1). Chr7 trisomy (Tri7) strains from patient MN were attenuated for hyphal and biofilm formation in vitro compared to euploid strains, due at least in part to *NRG1* over-expression. Nevertheless, representative Tri7 strain M1 underwent filamentation during disseminated candidiasis (DC) in mice. M1 was more fit than euploid strain M2 during DC and mouse gastrointestinal colonization, and in blood ex vivo. M1 and M2 exhibited identical echinocandin minimum inhibitory concentrations, but M2 was more tolerant to micafungin in vitro. Furthermore, M2 was more competitive with M1 in mouse kidneys following micafungin treatment than it was in absence of micafungin. Tri7 strains represented 74% of patient MN’s baseline BC population, but after 1d and 3d of echinocandin treatment, euploid strains were 93% and 98% of the BC population, respectively. Findings suggest that echinocandin tolerant, euploid strains were a subpopulation to more virulent Tri7 strains at baseline and then were selected upon echinocandin exposure. In conclusion, BCs in at least some patients are comprised of diverse *C. albicans* populations not recognized by the clinical lab, rather than single strains. Clinical relevance of *C. albicans* diversity and antifungal tolerance merits further investigation.

## Introduction

Clinical strains of *Candida* spp. from sites of mucosal colonization and longitudinal *Candida* strains from sites of invasive disease often demonstrate within-host genetic and phenotypic diversity.^1,2,3–6^ ^4,5,7^ The long-standing paradigm is that most fungal or bacterial sterile site infections reflect proliferation of a single, genetically identical strain that passes through a bottleneck (“single organism” hypothesis).^8–10^ In recent studies, however, we and others have shown that blood cultures (BCs) from patients with *Staphylococcus aureus*,□ carbapenem-resistant *Klebsiella pneumoniae* (CRKP) or *Candida glabrata* bloodstream infections (BSIs) can be comprised of genetically and phenotypically diverse strains, including strains unrecognized by the clinical laboratory that differ in antimicrobial susceptibility.^11–13^ Within-patient *C. glabrata* strains differed by single nucleotide polymorphisms (SNPs) and, to a lesser extent, insertions-deletions (indels), gene copy number variations, presence/absence of specific genes and chromosomal rearrangements.

*C. albicans* is the most intrinsically virulent *Candida* sp. and the leading cause of candidemia globally.^14^ *C. albicans* differs from *C. glabrata* in having a diploid, rather than haploid genome. In this study, we determined whether contemporaneous *C. albicans* strains from positive BCs of patients with BSIs were genetically and phenotypically distinct, and we compared diversity of strains from longitudinal BCs of patients with persistent BSIs.

## Results

### Genetic diversity of *C. albicans* strains

We performed whole genome sequencing (WGSing; Illumina NextSeq) on 10 strains from individual colonies in each of 4 patients [Supplemental Table 1]. For patient G, 10 strains were sequenced from the baseline (day (d)-0) BC, including the index strain from the clinical laboratory and a strain from each of 9 randomly-selected colonies. For other patients, index and 4 randomly-selected strains were sequenced from both baseline and last positive BCs (collected 3d (patient AB), 12 d (QR) and 13 d (MN) after baseline). Strains had 8 pairs of chromosomes and 14.24-14.82 Mbp genomes. On average, 98% of reads aligned to the reference *C. albicans* SC5314 genome (range: 96%-99%). Strains clustered by patient upon SNP phylogenetic analysis [Figure 1]. Within-patient strains differed by 0-7 (AB), 7-42 (G), 3-36 (MN) and 8-28 (QR) SNPs [Supplemental Table 2]. Within 3 patients, strains also differed by 1-8 (G), 0-9 (MN) and 1-6 (QR) indels.

**Figure 1.**
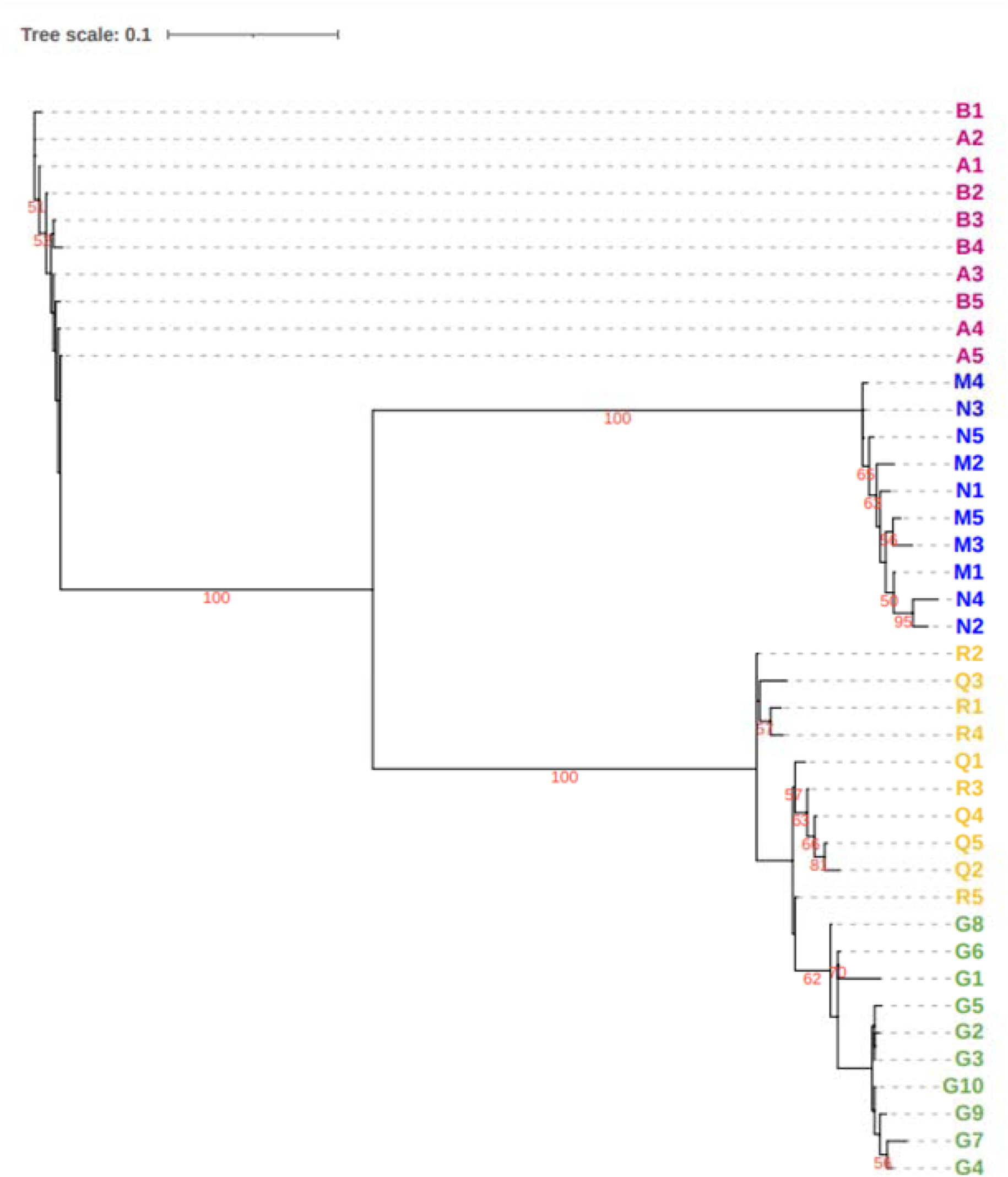
Core genome single nucleotide polymorphism (SNP) phylogeny of *Candida albicans* strains from patients with bloodstream infections. Ten strains from each of 4 patients (AB (magenta), MN (blue), QR (yellow) and G (green) underwent whole genome sequencing. The phylogenetic tree was built using RaxML under the general time-reversible model with 1,000-bootstrap replicates. Within-patient strains had the following ranges of SNP differences: patient AB (range, 0-7); patient MN (range, 3-36); patient QR (range, 8-28); patient G (range, 7-42). Nodes grouping by clade and patient had 100% bootstrap support values.

Large-scale, within-patient genetic differences among strains were observed in 3 of 4 patients (G, MN, QR) [Figure 2. Within 2 patients (G, MN), strains differed by whole chromosome aneuploidies. In patient G, index strain G1 exhibited trisomy of chromosome (Chr) 5 (Tri5); other strains were euploid [Figure 2A]. In patient MN, 4 of 5 strains from the baseline BC (index strain M1, M3-M5) exhibited trisomy of Chr7 (Tri7); 1 of 5 strains from baseline (M2) and all 5 strains from d13 BCs (index strain N1, N2-N5) were euploid [Figure 2B]. Ploidy was confirmed by quantitative real-time PCR (qPCR) [Supplemental Table 3; Supplemental Figure 1].^12,15^ In patient QR, loss of heterozygosity (LOH) of Chr1 was detected in 7 of 10 strains, including 2 of 5 baseline (index strain Q1 and Q2) and all 5 d12 (index strain R1, R2-R5) strains [Figure 2C]. R1-R5 also had segmental LOH of Chr7, which was not present in Q1-Q5. Within-patient AB strains did not show large-scale genetic variations.

**Figure 2.**
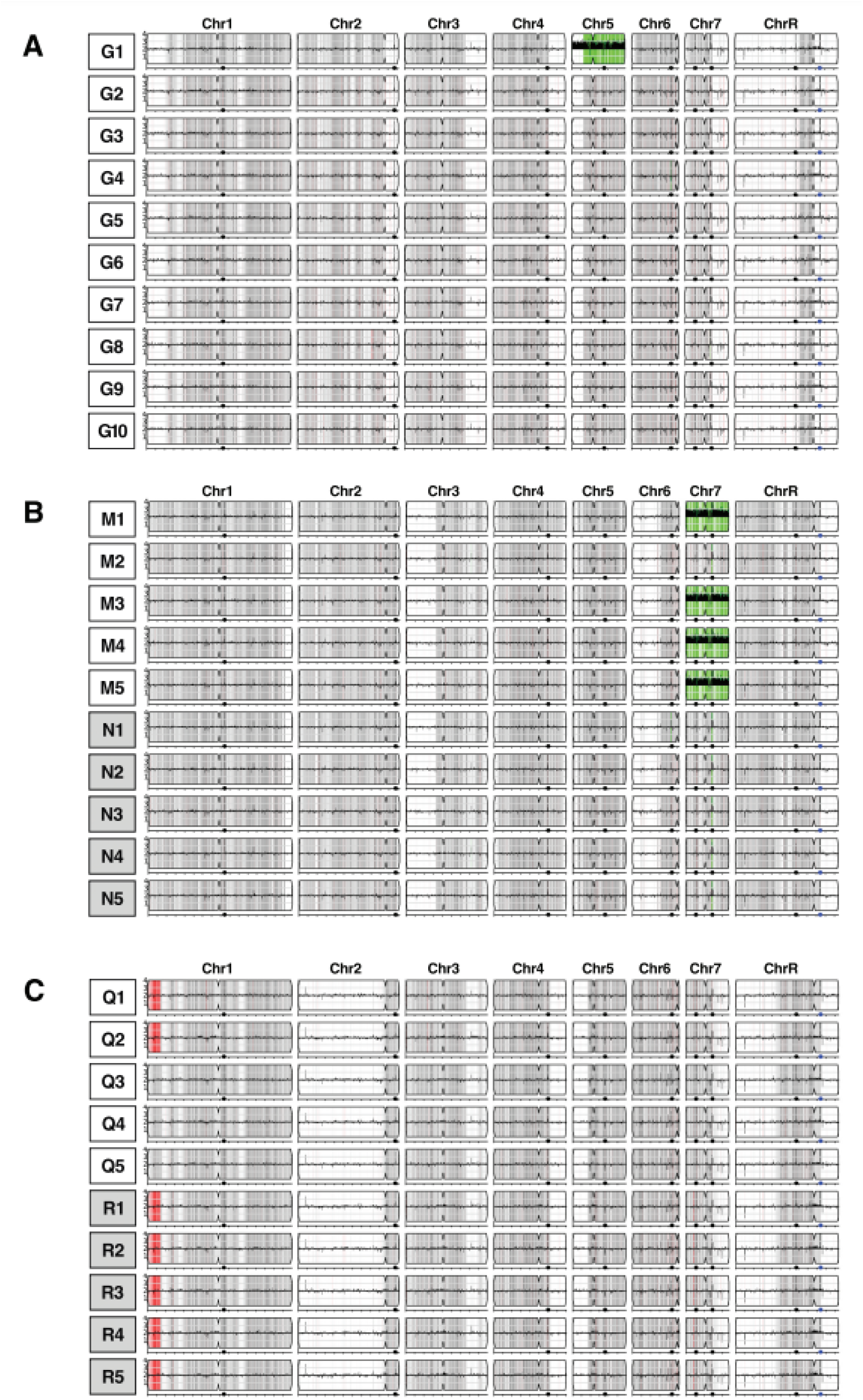
Copy number variants (CNVs), single nucleotide polymorphism (SNP) density and loss of heterozygosity (LOH) in *Candida albicans* strains from patients with bloodstream infections. Strains from patients G, MN and QR are shown in Figures A through C, respectively. Chromosomes are arranged along the horizontal (x) axis. The vertical (y) axis represents sequence read depth, scaled to actual copy number. CNVs are presented as black lines drawn upwards or downwards from the mid-line of the chromosome. Regions with CNV are highlighted in green. Regions of SNP density are presented as shades of grey, and homozygous regions are white. Regions of LOH are red. Black circles along the bottom of chromosomes represent major repeat sequences (MRSs). **A.** Strain G1 differed from other G strains by having an extra copy of whole Chr5 (i.e., trisomy 5), as evident by a block of black lines extending upward from the mid-line (green highlight). Strains did not differ by SNP density or LOH. **B.** Strains M1, M3, M4 and M5 differed from other MN strains by having an extra copy of whole Chr7 (i.e., trisomy 7). All MN strains had an extra copy of a small region adjacent to the second MRS of Chr7 (green highlight). Strains did not differ by SNP density or LOH. **C.** Strains Q1, Q2 and R1-R5 had a region of LOH (red highlight) in a segment at the end of the left arm of chromosome 1, which was not apparent in strains Q3-Q5. As indicated by grey highlight, the latter strains did not have gaps in SNP density in the region of LOH. A small potential region of LOH was also present uniquely in strains R1-R5, demarcated by a single red line near the middle of the left arm of Chr7; the size of this region was at or very near the resolution limit for YMAP analysis (4.5 Kb). QR strains did not differ by CNV or ploidy. The only differences among strains in SNP density was in the region of LOH.

### Screening phenotypes of *C. albicans* strains

All 40 strains were susceptible to echinocandins and azoles. Within-patient minimum inhibitory concentrations (MICs) were within 2-fold. Strains from patients AB and QR formed normal hyphae in liquid media (YPD with 10% fetal bovine serum, RPMI1640 with 2% glucose, Spider; 37°C) and on solid agar (RPMI1640 with 2% glucose, Spider, M199; 30°C). Strains from patient G were impaired in hyphal formation in liquid media and on solid agar [Supplemental Figure 2]. Strains from patient MN formed normal hyphae in liquid media. On solid agar, however, all 4 Tri7 strains (baseline strains M1, M3-M5) were impaired in hyphal formation, whereas all 6 euploid strains (baseline M2 and d13 N1-N5) formed robust hyphae [Figure 3]. Tri7 strains also produced <30% of biofilm that was generated by euploid strains, as measured by crystal violet assay [Figure 4; *p*-values< 0.0001].

**Figure 3.**
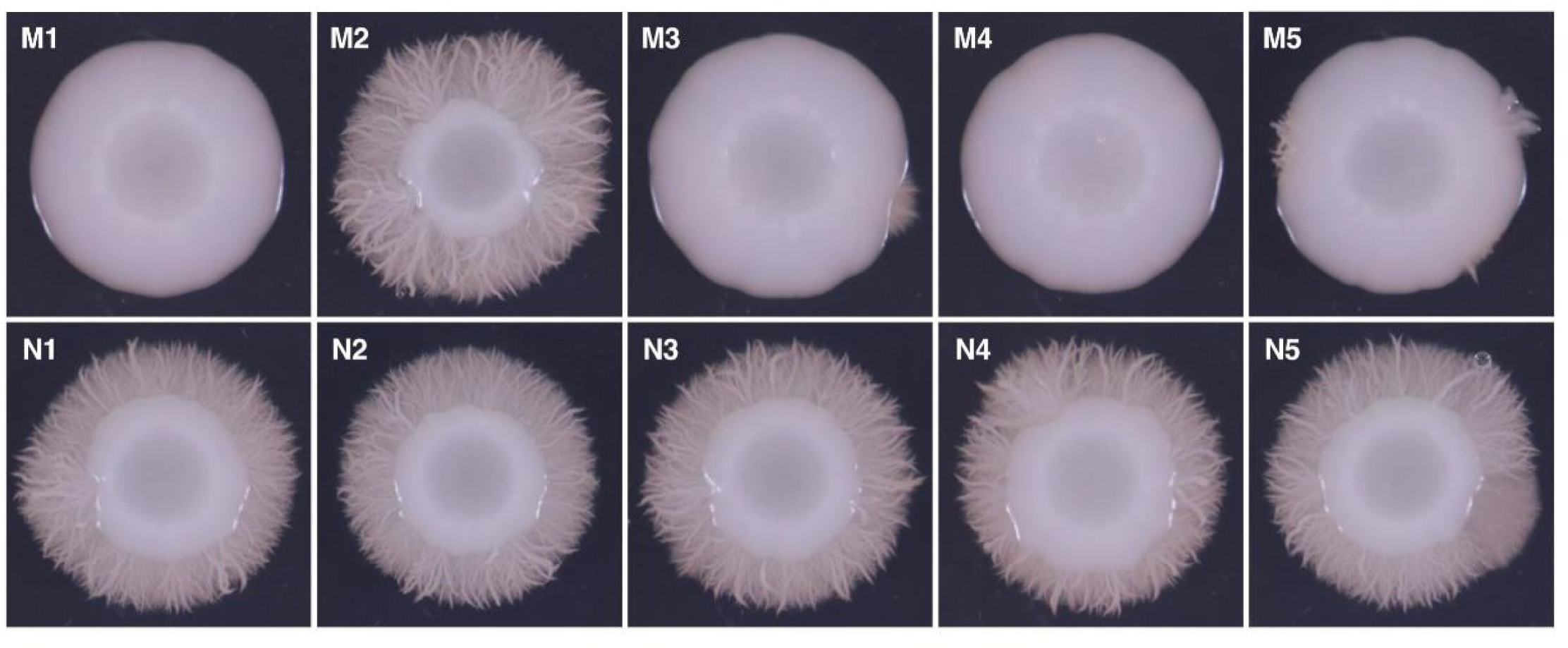
Hyphal formation by MN strains on solid agar. Strains were photographed after incubation on RPMI 1640 with 2% glucose medium at 30°C for 5 days. Euploid strains (M2, N1, N2, N3, N4 and N5) formed extensive hyphae, whereas trisomy 7 strains M1, M3, M4 and M5 did not.

**Figure 4.**
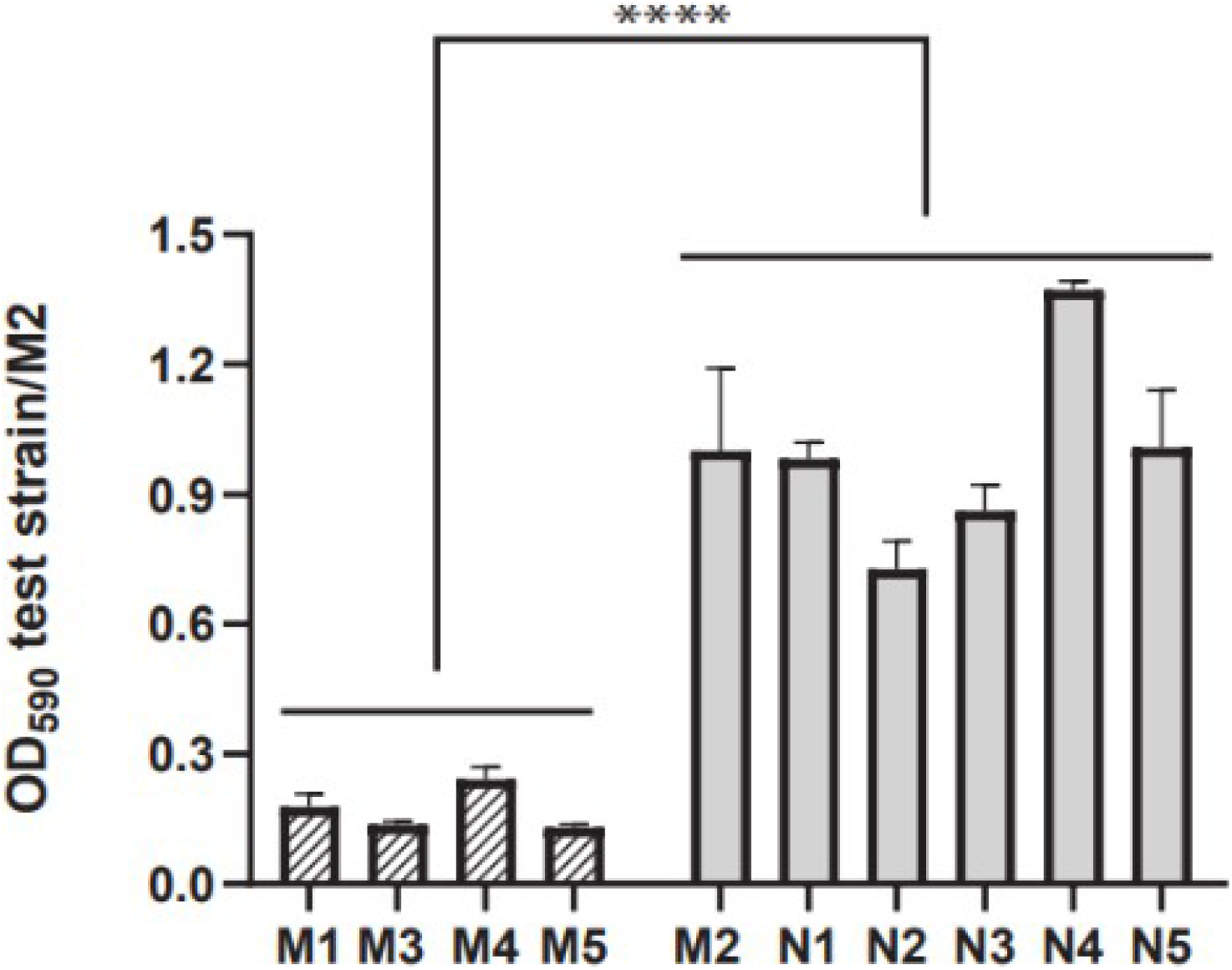
Biofilm formation by MN strains. Biofilm formation by MN strains at 37°C in RPMI1640 medium supplemented with 2% (w/v) glucose was measured by a 0.1% aqueous crystal violet assay in flat bottom wells as optical density at 590 nm (OD_590_). Data are presented as mean ± SEM of ratio of OD_590_ for a given strain relative to that of euploid strain M2 (3 independent experiments). Tri7 (M1, M3, M4 and M5) and euploid (M2, N1-N5) strains are represented by striped and solid grey bar graphs, respectively. Tri7 strains consistently produced < 30% of biofilm produced by euploid strains (mean ± SEM ratio: 0.17±0.01 *vs*. 0.99±0.04; \*\*\*\**p*<0.0001, Mann-Whitney test). Tri7 strains did not differ significantly from one another in biofilm formation, nor did euploid strains significantly differ among themselves.

### Phenotypic diversity of *C. albicans* strains from patient MN

We chose strains from patient MN for more in-depth phenotypic assays because they demonstrated genetic differences that correlated with differences in hyphal and biofilm formation, and serial positive BCs over 13d were available for study. The timeline of *C. albicans*-positive BCs and antifungal treatment is shown in Supplemental Figure 3.

#### Micafungin responsiveness

We first tested serial ten-fold dilutions of each strain for responsiveness to micafungin and other stress-inducing agents in dot-blot assays. Tri7 strains (M1, M3, M4, M5) were more susceptible to Congo red (200 µg/mL), sodium dodecyl sulfate (0.04%) and micafungin (0.03 µg/mL) at 30°C and 39°C than were euploid strains [Figure 5A]. There were no significant differences among strains in presence of fluconazole (1 µg/mL), methyl methane sulfonate (0.02%), hydrogen peroxide (H_2_O_2,_ 2.5mM), sodium chloride (1M) or caffeine (10 mg/mL).

**Figure 5.**
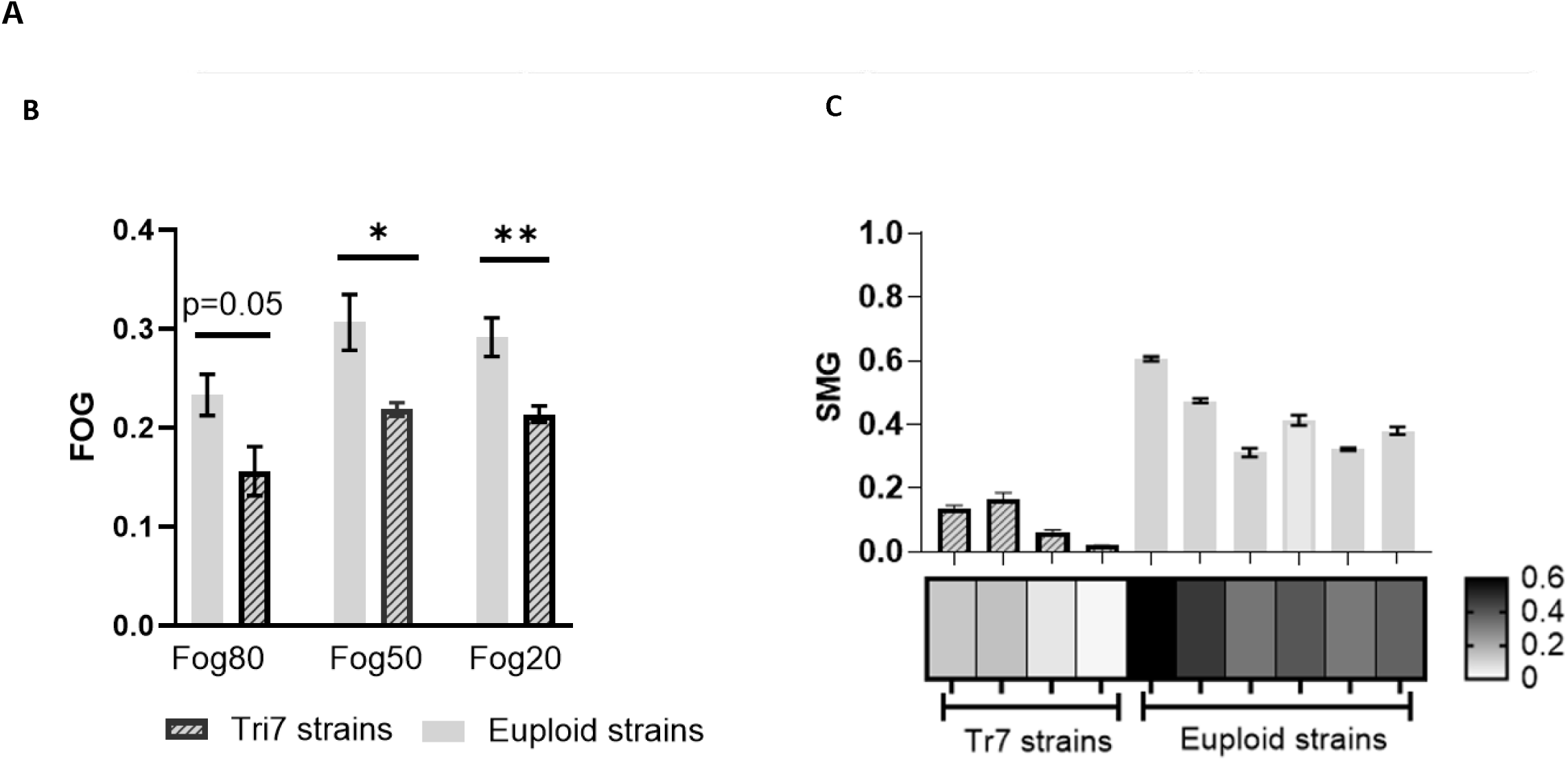
Susceptibility of MN strains to cell wall-active agents and tolerance to micafungin. **A.** In susceptibility experiments, serial 10-fold dilutions starting at 1×10^6^ CFU/mL of trisomy7 and euploid strains were spotted at exponential growth onto Sabouraud dextrose agar (SDA, control) and SDA containing Congo red (200μg/mL), sodium dodecyl sulfate (SDS, 0.04%) or micafungin (0.03 μg/mL). Plates were incubated at 30°C or 39°C for 3d. Images are shown for representative trisomy7 (M1, M3) and euploid (M2, N1) strains on d3 at 39°C. Phenotypes were similar at 30°C. **B.** Micafungin tolerance was first assessed using a disk diffusion assay to measure fraction of growth (FoG). A filter disc inoculated with 5 µg of micafungin was placed in the middle of a casitone agar place saturated with *C. albicans* strains. Tolerance was determined at 48h growth at 35°C. FoG was defined as growth within the region of inhibition relative to the maximum growth. FoG was calculated by diskImageR and image J at 80%, 50% and 20% relative inhibition (FOG_80_, FOG_50_ and FOG_20_, respectively). Data shown are mean ± standard error of means of FoG values for Tri7 and euploid strains. *P*-values were determined by student’s t test with Welsh correction (* denotes p <0.05 and ** p<0.01). **C.** Micafungin tolerance was also assessed using a broth microdilution assay to measure supra-MIC growth (SMG, defined as the proportion of average of growth at 48h in wells with drug concentration above the MIC relative to the growth of no drug control wells). SMG data (OD_600_ at 48h) are presented as mean ± standard error of euploid and Tri7 strains on a bar graph (top) and as a heat map (bottom). Heat map scale is shown on the bottom right.

We next evaluated MN strains for micafungin tolerance (enhanced growth in presence of drug compared to control, without changes in MIC) using disk diffusion (fraction of growth (FoG)) and liquid microdilution (supra-MIC growth (SMG)) assays.^16,17^ FoG of euploid strains within zones of 80%, 50% and 20% inhibition by micafungin (FoG_80_, FoG_50_, FoG_20_) were higher than those of Tri7 strains (*p*=0.05, 0.03 and 0.008, respectively) [Figure 5B]. Likewise, mean SMG (average growth within micafungin-containing wells above MIC_50_/growth in absence of micafungin) of euploid strains was higher than that of Tri7 strains (*p*=0.0003) [Figure 5C]. Therefore, euploid strains were more tolerant than Tri7 strains to micafungin by both FoG and SMG.

#### *NRG1* dose effect

Decreased filamentation of some *C. albicans* Tri7 strains has been linked to dose-dependent expression of Chr7 gene *NRG1*, which encodes a negative regulator of certain hyphal genes.^15^ We demonstrated by qRT-PCR that *NRG1* expression by Tri7 strain M1 was ∼1.5 fold higher than that by euploid strains [Supplemental Figure 4A]. ^18^ We created *NRG1* over-expression strains in euploid M2 and N1 backgrounds (strains M2-Nrg1OE and N1-Nrg1OE, respectively). *NRG1* expression by M2-Nrg1OE and N1-Nrg1OE was ∼1.8-fold and ∼1.6-fold higher than by the respective euploid parent strains (*p*=0.001) [Supplemental Figure 4A]. Over-expression strains had reduced hyphal and biofilm formation compared to euploid parent strains (biofilm *p*-values <0.0001) [Supplemental Table 3; Supplemental Figures 4B-C].

#### Competitive growth in vitro and in blood ex vivo

Since Tri7 and euploid strains comprised a mixed population within patient MN’s baseline BC, we studied competitive fitness of bar-coded Tri7 strain M1 and bar-coded euploid strain M2 under various conditions. Bar-coded strains did not differ from respective parent strains during growth in YPD media or by echinocandin MICs or in hyphal and biofilm formation.

We conducted competitive growth assays (1:1 ratio, 1×10^4^ colony forming units (CFUs)/mL each) in liquid YPD media at 39°C in presence or absence of H_2_O_2_ (1 mM) or micafungin (0.125 μg/mL).^15^ There were no significant differences in competitive fitness between strains over 3, 6 or 9d in YPD alone or in presence of H_2_O_2_, as determined by bar code sequencing [Figure 6A]. Strain M2 was more fit than M1 at each time point in the presence of micafungin (*p* <0.0001), consistent with FoG and SMG tolerance data. During ex vivo competitive assays in fresh blood from a healthy volunteer at 37°C (1×10^4^ CFUs)/mL each), M1 was significantly more fit than M2 over 1, 3 and 7d [Figure 6B].

**Figure 6.**
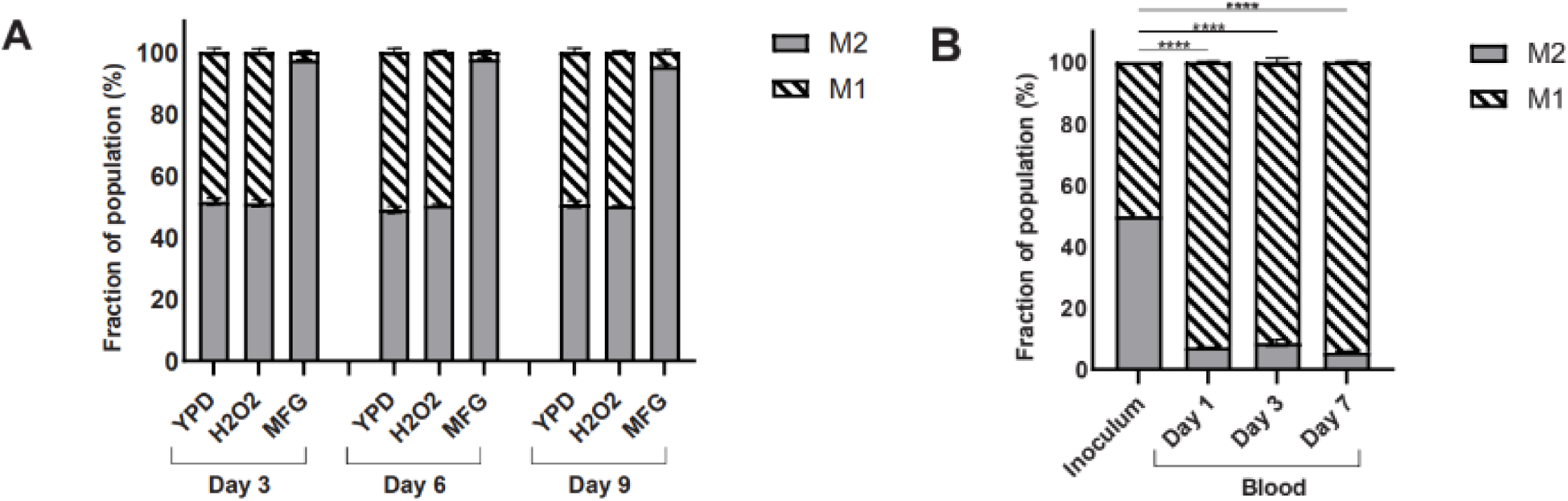
Competitive fitness of trisomy 7 strain M1 and euploid strain M2 in vitro and ex vivo. Barcoded strains M1 and M2 were mixed 1:1 in all experiments; input ratios were confirmed by colony forming unit determinations. At given timepoints, competitive indices (fraction of total population) of M1 and M2 were determined by barcode sequencing. Data are presented as mean ± SEM competitive indices for each strain from 3 independent experiments. * *p*<0.05, ** *p*<0.01, \*\*\**p*<0.001, \*\*\*\**p*<0.0001 **A. In vitro**. Barcoded M1 and M2 were co-cultured in yeast peptone dextrose (YPD) medium at 39°C with or without H_2_O_2_ (1 mM) or micafungin (0.125 µg/mL) for 3, 6 and 9d. Strains did not differ in fitness in YPD alone or in presence of H_2_O_2_. M1 was significantly more fit than M2 at each time point in presence of micafungin (*p*< 0.0001; unpaired student’s t test). **B. Ex vivo.** Strains were co-incubated in blood from a healthy human volunteer at 37°C. Fractions of each strain out of the total population were assessed at d1, d3 and d7. Strain M1 was significantly more fit than M2 at each time point. *P*-values were calculated using one-way ANOVA with Dunnett’s multiple comparisons against inoculum fraction.

#### Competitive mouse models of GI colonization and disseminated candidiasis (DC).^15^

We co-infected male and female mice with bar-coded M1 and M2 (1×10^8^ CFUs each) by gavage. M1 significantly out-competed M2 within stool on d1, d3 and d7 [Figure 7A], and in ileum and colon on d7 [Figure 7B]. We then co-infected mice via lateral tail vein with bar-coded strains (5×10^5^ CFUs each). M1 significantly out competed M2 within kidneys, the major DC target organ, at d1, 3 and 7 [Figure 7C]. We also treated a group of M1:M2 co-infected mice with a single dose of micafungin (8.3 mg/kg/mouse intraperitoneal, 30 minutes post-infection). Strain M2 was 59% more competitive with M1 in kidneys at 24h following treatment than it was in absence of micafungin [Figure 7C].

**Figure 7.**
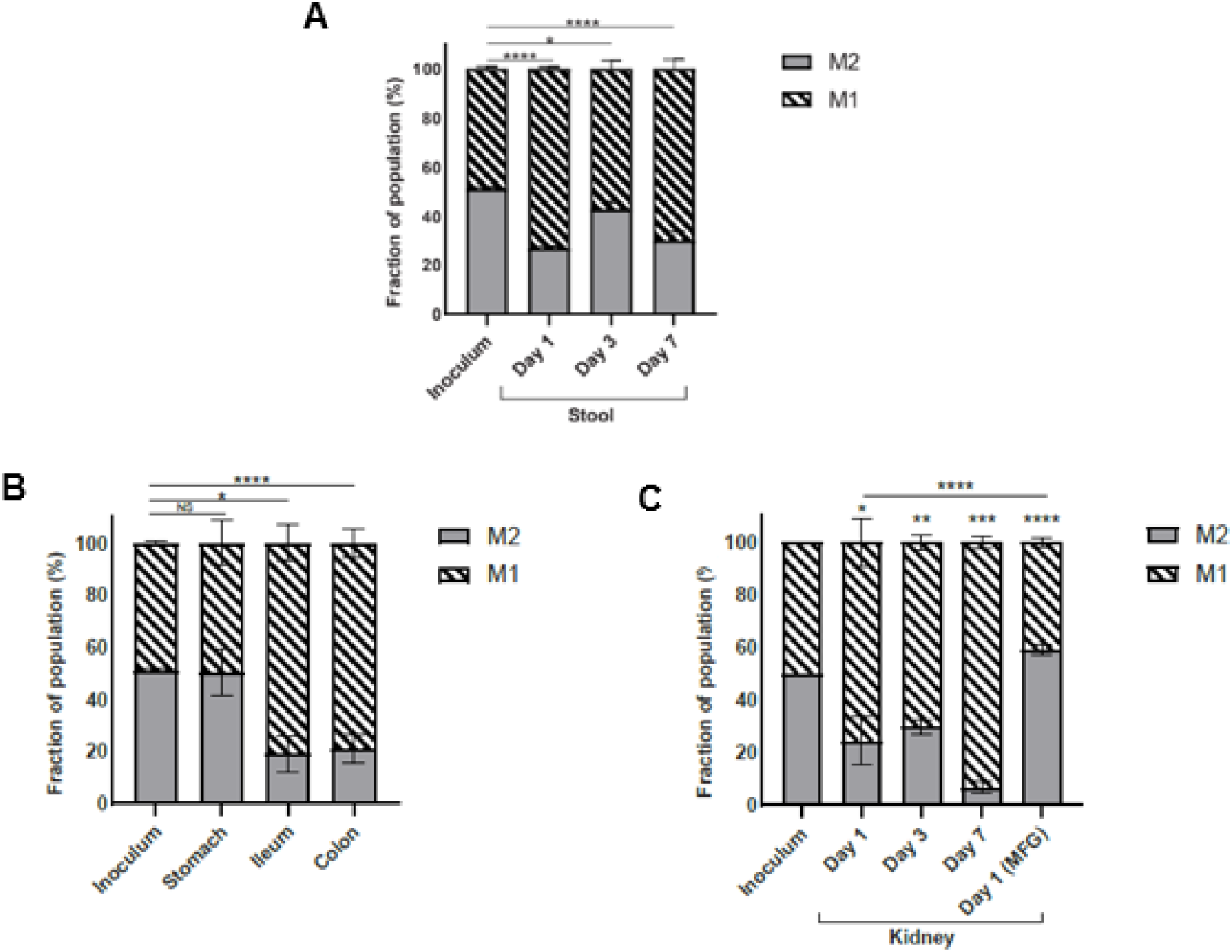
Competitive fitness of trisomy 7 strain M1 and euploid strain M2 in mouse models. Barcoded strains M1 and M2 were mixed 1:1 in all experiments; input ratios were confirmed by colony forming unit determinations. At given timepoints, competitive indices (fraction of total population) of M1 and M2 were determined by barcode sequencing. Data are presented as mean ± SEM competitive indices for each strain from 3 independent experiments. * *p*<0.05, ** *p*<0.01, \*\*\**p*<0.001, \*\*\*\**p*<0.0001 **A, B. Mouse model of gastrointestinal (GI) colonization**. Mice were co-infected via gavage with barcoded M1 and M2 (1×10^8^ CFU of each strain per mouse). Stool samples were obtained on d1, d3 and d7 after inoculation (Figure 7A). On d7, mice were sacrificed, and stomach, ileum and colon were obtained and homogenized for DNA extraction for barcode sequencing (Figure 7B). M1 was significantly more fit than M2 within stool at all time points, and more fit than M2 within ileum and colon. Strains did not differ in fitness within stomachs. **C. Mouse model of disseminated candidiasis (DC).** Mice were co-infected via lateral tail vein with barcoded strains M1 and M2 (5×10^5^ CFU of each strain per mouse). Kidneys were obtained and homogenized for DNA extraction for barcode sequencing on d1, d3 and d7. A group of co-infected mice were treated with a single dose of micafungin (8.3 mg/kg/mouse intraperitoneal, 30 minutes post-infection), and kidneys were obtained for barcode sequencing 24h following treatment. *P*-values for were determined using one-way ANOVA test with Dunn’s multiple comparison against inoculum fraction. Tri7 M1 strain was more fit than M2 in the GI and DC models. M2 was significantly more fit within kidneys of mice treated with micafungin than it was within kidneys of untreated mice. NS: non-significant; MFG: micafungin

#### Mono-infection DC

Finally, we performed single strain infections in which mice were inoculated intravenously with either bar-coded M1 or M2 (1×10^6^ CFU). M1 burdens were significantly higher than M2 burdens within kidneys at d1 and 3 [Supplemental Figure 5]. Both strains were mixtures of yeast and filamentous morphologies within kidneys [Supplemental Figure 6].

### Tri7 strains in longitudinal and spiked BCs

Echinocandin treatment in patient MN was initiated on d2. To estimate point prevalence, we screened 96 strains from longitudinal BCs for attenuated biofilm formation (< 30% of that formed by euploid strain M2) as marker for Tri7 and confirmed results for certain strains by qPCR. Percentages of Tri7 strains were 74% (baseline, 71/96), 7% (d3, 7/96), 2% (d5, 2/96), 2% (d10, 2/96) and 0% (d13, 0/96) [Supplemental Figure 3].

To assess Tri7 stability, we spiked M1 into a sterile BC bottle and incubated at 37°C until turbidity was evident (24-48h). Aliquots were sub-cultured on SDA plates at 30°C for 48h, and each of 10 strains from randomly selected colonies was shown to still harbor Tri7 by WGS [Supplemental Figure 7].

## Discussion

We demonstrated that *C. albicans* strains from BCs of individual patients were genetically and phenotypically diverse. The most striking within-patient genetic differences among strains were aneuploidies and LOH, which were identified in 3 of 4 patients. In patient MN, Tri7 and euploid strains were equally fit in liquid media, and they exhibited similar micafungin MICs. Euploid strains, however, were more tolerant to micafungin than Tri7 strains by FoG, SMG and competitive growth assays. In contrast, Tri7 strain M1 was more fit than euploid strain M2 during growth in human blood ex vivo, and during mouse models of GI colonization and DC. M2 was more competitive than M1 in mouse kidneys following micafungin treatment of DC than it was in the absence of micafungin. Tri7 strains predominated in patient MN’s baseline BC (74% of the population), but they were largely replaced by euploid strains after 1 and 3d of echinocandin treatment (93% and 98% euploid, respectively). By d13 of BSI, 100% of BC strains were euploid. The data suggest that Tri7 strains were more virulent than euploid strains, which conferred an advantage at baseline in absence of antifungal treatment. Echinocandin tolerant, euploid strains that were present at baseline but unrecognized by the clinical lab emerged as dominant upon echinocandin exposure. These results and our previous findings for *C. glabrata*- and CRKP-positive BCs challenge the single organism hypothesis of BSI pathogenesis, and they support a population-based paradigm for BSIs by commensal microbes in at least some patients.^12,13^

Large-scale genetic variants like aneuploidy and LOH, including Tri7, are described in clinical *C. albicans* strains and in strains exposed to antifungals in vitro or recovered from mouse models of candidiasis.^5,15,19–22^ Aneuploidy can involve any *C. albicans* Chr, with positive or negative consequences for fitness and antifungal susceptibility depending on specific Chr, strain background and environmental conditions.^23–25^ Findings that Tri7 strain M1 was attenuated for hyphal formation in vitro compared to euploid partner strain M2, while exhibiting enhanced GI colonization, invasion of ileum and colon, and hematogenous infection of kidneys in mice, were consistent with data for certain previously studied Tri7 strains.^15,22,25^ Echinocandin tolerance has been described for *C. albicans* strains with Chr2 and Chr5 aneuploidies, but it has not been associated with Tri7.^26–30^ Fluconazole tolerance has been reported in some *C. albicans* strains causing persistent BSIs despite treatment,^16^ but neither fluconazole nor echinocandin tolerance are broadly validated as determinants of outcomes in infected patients.^16^ ^26,30^ We did not identify echinocandin-resistant *C. albicans* strains. In our *C. glabrata* study, baseline BCs in 2 of 10 patients were a mixture of fluconazole-susceptible and originally unrecognized fluconazole-resistant strains, the latter of which were later recovered as index strains from recurrent infections. Recently, echinocandin heteroresistance (a low-frequency subpopulation of resistant cells) among *C. parapsilosis* was connected to echinocandin break-through BSIs.^31^ There is pressing need for multicenter studies of the clinical relevance of phenotypes like antifungal tolerance and heteroresistance that are not detected by standard clinical laboratory susceptibility testing methods.

As is true for antifungal tolerance, the clinical relevance of diversity in virulence among *Candida* strains is uncertain. While hyphal and biofilm formation are classic *C. albicans* virulence determinants, it is more precise to understand bidirectional regulation of morphogenesis and biofilm maturation, as dictated by local environment, as crucial to commensalism and pathogenesis.^32^ It is well recognized that certain *C. albicans* strains, like Tri7 strains here, manifest filamentation defects on solid agar, but not in liquid media or within target organs. Moreover, strains with attenuated filamentation or biofilm formation in vitro may manifest full or even heightened pathogenic capacity in vivo. In the end, *C. albicans* is an opportunistic pathogen that does not depend upon a dominant virulence factor, but rather a complex interplay of properties that optimize survival in various in vivo niches.^14^ At least in part, attenuated hyphal formation by Tri7 strains in vitro has been attributed here and elsewhere to over-expression of *NRG1*, which was initially described as a transcriptional repressor of *C. albicans* SC5314 hyphal and stress response genes.^15^ It is now apparent that *NRG1* exerts variable effects on gene expression, filamentation, fitness and virulence depending on strain background.^15,33,34^ Tri7 was not previously linked to aberrant *C. albicans* biofilm formation, although high *NRG1* expression was shown to limit biofilm maturation and promote dispersion of yeast cells from immature biofilm.^35^

Tri7 was stable in strain M1 following incubation in spiked BC bottles, suggesting that euploid strains in patient MN emerged in vivo rather than ex vivo. This finding is notable since changes in *Candida* ploidy are often reversible adaptations to stress.^36^ We propose that most of the *C. albicans* and *C. glabrata* strain diversity we observed in BCs was likely to have arisen during GI commensalism (or at another colonization site).^6^ We recognize that we may have failed to identify important genetic or phenotypic variants by studying only 10 *Candida* strains per patient. It is feasible that *C. albicans* genetic variants other than differences in Chr7 ploidy also contribute to important phenotypes. In our previous study, the predominant within-patient genetic differences among *C. glabrata* strains from baseline BCs of 10 patients were core genome SNPs and indels.^12^ Within-patient SNPs and indels were less prevalent among *C. albicans* strains from BCs of 4 patients. Future studies will elaborate whether differences between spp. have biologic explanations, reflect features that are particular to individual cases, and/or stem from relatively small sample sizes. To date, our studies are the only investigations of within-patient diversity among *Candida* strains during BSIs. Results cannot necessarily to extrapolated to other *Candida* spp., other opportunistic commensal microbes, or other sites of infection.

In conclusion, positive BCs contained mixed populations of *C. albicans* strains that differed most strikingly by aneuploidies and LOH, and in echinocandin tolerance and virulence attributes. Studies to assess the clinical relevance of *Candida* diversity are indicated, as is research into understanding how diversity and adaptation of various microbes facilitates commensalism and disease. If results here and from our studies of *C. glabrata* and CRKP BSIs are validated, clinical and microbiology lab practices may need to be revised to consider microbial populations.

## Methods

This study was approved by the University of Pittsburgh Institutional Review Board through Expedited Review according to OHRP, 45 CFR 46.110 and FDA 21 CFR 56.110, and by the University of Pittsburgh Animal Care and Use Committee. Reagents, primers and plasmids appear in Supplemental Tables 3, 4 and 5, respectively. BC bottles and index strains from patients with *C. albicans* BSI were obtained from the UPMC clinical microbiology laboratory.^12^ We streaked 10□µL from bottles onto 2 SDA plates and incubated for 48h at 35°C. For each BC, a strain isolated from each of 4–9 morphologically indistinguishable colonies and the index strain underwent Illumina NextSeq WGSing [Supplemental Table 4].

### Illumina NextSeq and bioinformatic analysis

DNA extraction, library preparation and genome analysis were performed as previously published,^12^ and as described with details of variant filtration in Supplemental Methods.^37–41^ We selected gene variants called in ≥ 1 strain, but not all strains within each patient, focusing on non-synonymous substitutions, disrupted start/stop codons, and frameshift indels. Aneuploidy, gene deletions/duplications and LOH were analyzed using Ymap ^42^ and reference genome SC5314 (CGD: A21-s02-m09-r10). We used allelic ratio (homozygous: 0 or 1; heterozygous: 0.5) to determine whether a region with SNPs in the parent/reference strain had undergone LOH. Allelic ratio was calculated for each coordinate as: (number of reads with the more abundant base call)/(total number of reads). We selected internal strains for each patient to create a haplotype map and investigate within-patient LOH.

### In vitro phenotypes

Growth rate, filamentation, antifungal susceptibility, micafungin tolerance, dot blot and biofilm assays were performed using well-established protocols, as described in detail in Supplemental Methods.^16,17,43^

### Chromosome copy numbers

Chr5 and Chr7 copy numbers were determined by qPCR using multiple primer pairs, as previously described [Supplemental Table 5]. Chr1 and Chr4 qPCRs were copy number controls.^14^

### qRT-PCR transcriptional analysis

was performed using SYBR Green qPCR Master Mix.^15^ Expression was calculated using ΔΔCT, with normalization to housekeeping genes *ACT1* and 18S rDNA.

### *NRG1* over-expression strain

The *NRG1* promoter was replaced with the constitutively expressed *TDH3* promoter from plasmid pCJN542 (provided by Aaron Mitchell).^44^ Euploid strains M2 and N1 were transformed using PCR products from pCJN542 and primers NRG1-OE-F and NRG1-OE-R. Overexpression strains were selected on YPD+clonNAT (YPD, 200 µg/mL nourseothricin sulfate) plates.

### Barcoded strains

were constructed as previously described.^15^ Plasmids RB793 (Addgene #199121) and RB794 (Addgene #199122) containing barcodes were digested with NgoMIV and transformed into target strains. Transformants were selected on YPD+200 µg/mL nourseothricin. Integration at *NEUT5L* was confirmed by PCR using primers in Supplemental Table 4.

### In vitro competitive growth

Barcoded strains were grown overnight in liquid YPD at 30□C, washed with PBS and diluted in YPD. Strains (1:1 ratio, 1×10^4^ CFU/mL each) were incubated at 39°C with shaking (200rpm). Micafungin (0.125 μg/mL) or H_2_O_2_ (1 mM) were added on d0, 3 and 6. On d0, 3, 6 and 9, DNA was extracted from 200μL aliquots for bar code sequencing.

### GI colonization model

We used 8 mice at each time point (equal male:female, 4-6 weeks old, 20-25 g, ICR CD1, Envigo). GI infections were performed as previously described.^15^ Four days before infection, drinking water was supplemented with 1.5mg/mL ampicillin, 2mg/mL streptomycin and 2.5% glucose. Mice were gavaged with 1X10^8^ CFU of each strain. Fecal pellets from d1, 3, and 7 were homogenized in sterile PBS. On d7, stomach, ileum and colon were homogenized in PBS. Homogenates were plated on SDA plates with 200µg/mL each of ampicillin, streptomycin, chloramphenicol, kanamycin and nourseothricin, and processed for DNA extraction.^15^

### DC model

We intravenously inoculated 6 and 8 mice (equal male:female) per time point for co-infections and mono-infections, respectively; inocula were 5×10^5^ CFU per barcoded strain (co-infection) or 1×10^6^ CFU (mono-infection). In co-infections, an additional 6 mice were treated with micafungin (8.3 mg/kg/mouse intraperitoneal, 30 minutes post-infection) and sacrificed at 24h. Kidneys were homogenized for DNA extraction and barcode sequencing on d1, 3 and 7 post-infection (co-infection), or for tissue burden determination on SDA (mono-infection).^15^

### Barcode sequencing

DNA (1-3 μL) extracted from in vitro experiments, fecal samples and organ homogenates (Zymobiomics DNA Miniprep kit; Zymo Research) was used for PCR^14^ [Supplemental Table 4]. PCR parameters were 98°C for 30s, 98°C for 10s, 60°C for 30s, 72°C for 30s; steps 2 to 4 repeated for 34 cycles; and 72°C for 5 min. PCR products were gel purified (Wizard SV gel and PCR clean-up system, Promega). ExpressPlex library preparation and sequencing was done by SeqWell (MA, USA). Strains were identified by unique barcode using CLC Genomic workbench v.22.0.1. BC1/BC2 ratio was calculated and normalized by initial inoculum. BC2 percent was calculated as: (1/(Ratio+1))X100%.

### Populations in longitudinal BC bottles

Ninety-six strains from each BC in patient MN were screened for attenuated biofilm formation as a marker for Tri7 (mean biofilm OD_590_ of 3 independent experiments <30% of mean biofilm formed by euploid strain M2). Ploidy was confirmed by qPCR for ≥ 10 strains from each of d0 and d13 BCs.

### BC bottle spiking

A single colony of Tri7 M1 suspended in 50 µl saline was inoculated into a sterile BC bottle (Fisher Scientific BD BACTEC Plus Aerobic medium BD442023) that contained 5□mL blood from a healthy volunteer (∼1 CFU/mL), and incubated at 37°C with shaking (225rpm) until turbidity was evident.^12^ Ten microliters were streaked on an SDA plate, and incubated at 30°C for 48h. Single strains were isolated from 5-10 randomly chosen colonies for DNA extraction and Illumina sequencing.

### Statistical analyses

were performed using GraphPad Prism v9.1.2. Symmetric and asymmetric data are presented as means and standard error and as medians and interquartile ranges, respectively. CFU/g were log-transformed prior to analysis. Student t-test or Mann-Whitney U tests were used for comparisons of 2 groups, and one-way ANOVA or Kruskal-Wallis for comparisons of >2 groups. For all analyses, *p*<□0.05 (two-tailed) was significant.

## Data availability

Datasets are available in data files submitted with this article. WGS data have been deposited in the NCBI database under accession number PRJNA1118730. BioProject and associated SRA metadata are at: https://dataview.ncbi.nlm.nih.gov/object/PRJNA1118730?reviewer=nuvb3aprq91400u9pri6s57r97

## Acknowledgements

This project was supported by VA Merit Review 1IO1BX001955 (Dr. Clancy) and NIH R21 AI160098 (Dr. Nguyen) awards. Some of the data were presented at IDWeek 2024 (October 14, 2024, Los Angeles, CA). The authors do not report any conflicts of interest.

---

**Supplemental Figure 1.**
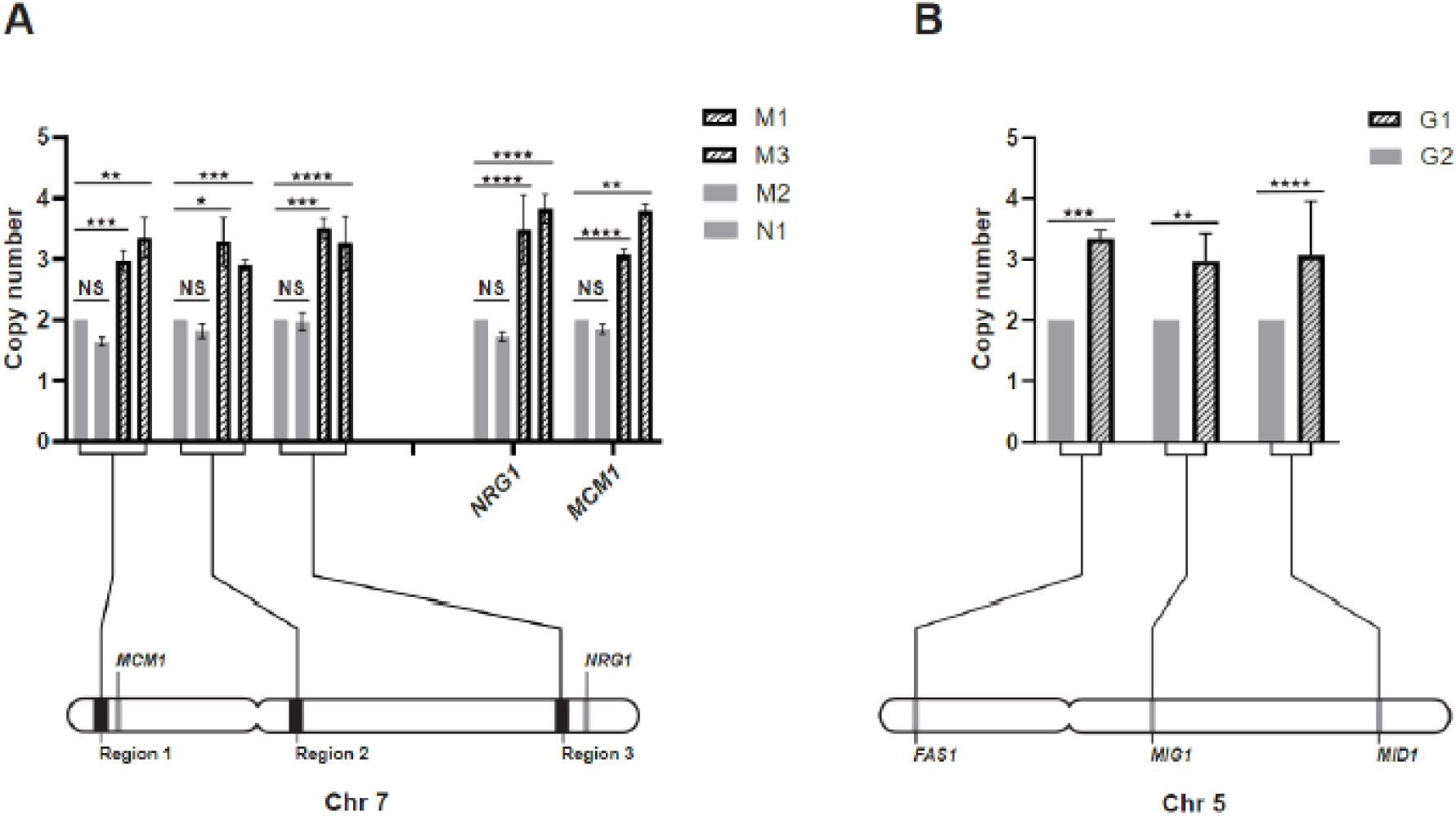
Validation of trisomy of chromosomes 7 and 5 by qPCR. Chr7 and 5 copy numbers were assessed in strains from patients MN and G, respectively, by qPCR. **A.** For patient MN, 3 target regions and 2 genes (*NRG1* and *MCM1)* on Chr7 were assessed for 2 euploid (M2, N1) and 2 Tri7 (M1, M3) strains. **B.** For patient G, genes *MID1*, *MIG1* and *FAS1* on Chr5 were assessed for a euploid (G2) and a Tri5 (G1) strain. Copy numbers for strains M2, N1 and N2 and strain G1 were calculated relative to those of strains M2 or G2, respectively. Target regions of Chr1 and Chr4 were used for normalization. Data are presented as means ± SEM from 3 independent experiments. Statistical analyses were performed using ANOVA with Dunnett’s (MN strains) or Sidak’s (G strains) multiple comparisons tests. *P*-values <0.05 were considered significant: \**p*-values <0.05, \*\**p*<0.01, \*\*\**p*<0.01, and \*\*\**p*<0.0001.

**Supplemental Figure 2.**
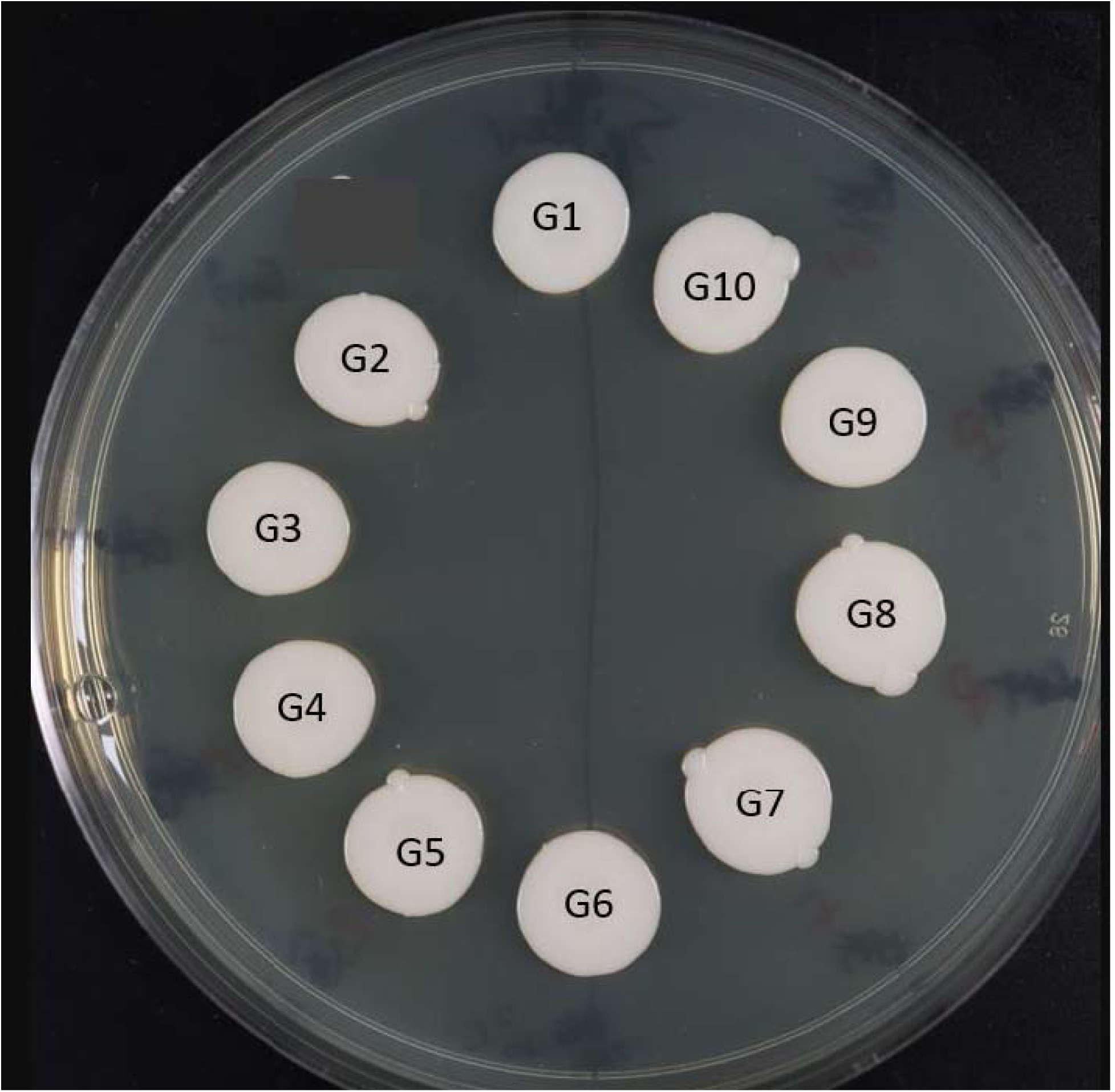
Impaired hyphal formation on Spider agar medium by strains from patient. **G.** Strains were grown under hyphal inducing conditions at 30°C. The picture here was taken after 14 days. Note lack of hyphal formation. Strains were also impaired in hyphal formation on other solid agar (M199, RPMI1640 at 30°C) and in liquid media (Yeast peptone dextrose with 10% fetal bovine serum; RPMI1640 at 37°C) (data not shown).

**Supplemental Figure 3.**
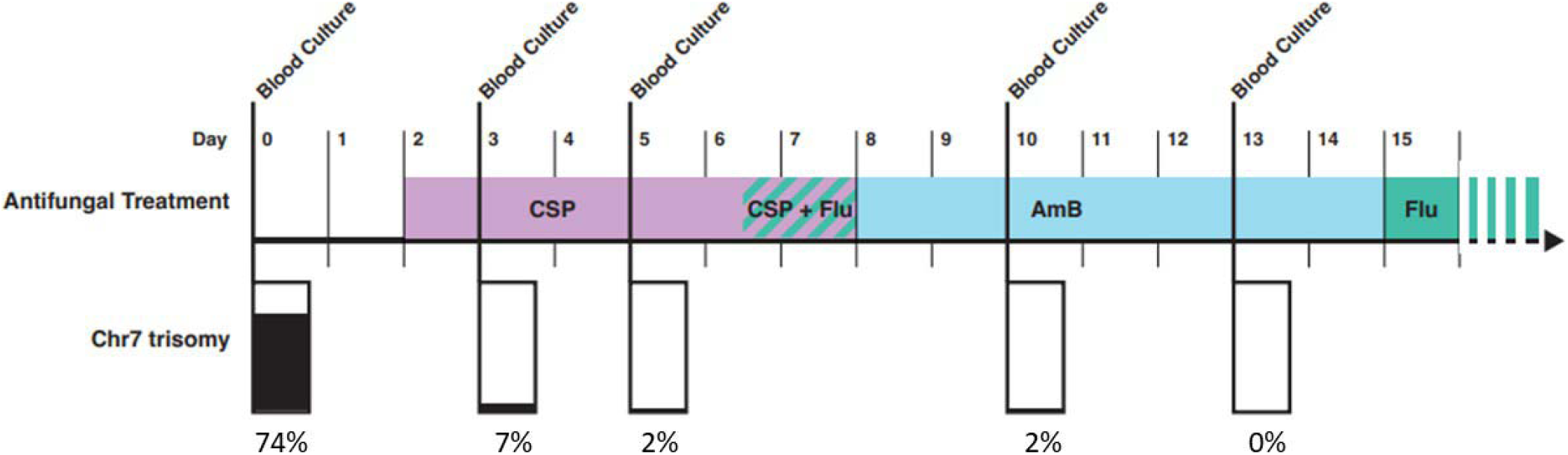
Timeline of *Candida albicans*-positive blood cultures (BCs) and antifungal treatment for patient MN. The baseline positive BC (i.e., BC at time of initial diagnosis) was collected on day 0. Antifungal treatment was initiated with caspofungin on day 2, after baseline BC results were reported by the clinical microbiology lab. Subsequent BCs that were positive for *C. albicans* were collected on days 3, 5, 10 and 13. The percentage of strains from each positive BC that had trisomy of Chr7 (Tri7) is shown at the bottom of the figure and represented as black in the bar graph. The percentage of strains that were euploid for Chr7 are represented by white in the bar graph. CSP, caspofungin; FLU, fluconazole; AmB, amphotericin B

**Supplemental Figure 4.**
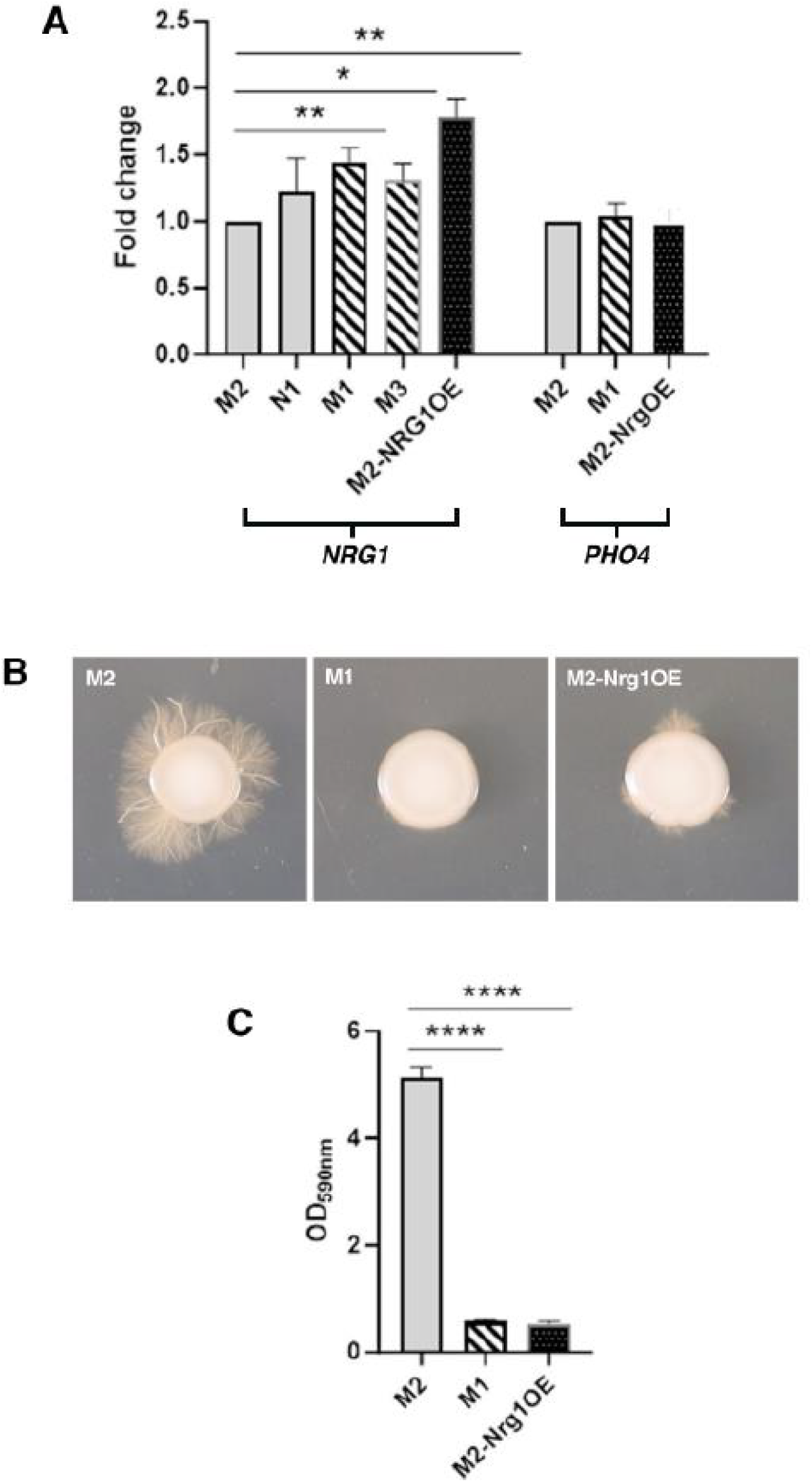
Impact of *NRG1* expression on biofilm and hyphal formation. **A.** *NRG1* expression by Tri7 strains (M1, M3), euploid strains (N1, M2) and M2-Nrg1OE (*NRG1* over-expression strain created in M2 background) was evaluated by qRT-PCR. Expression of *PHO4* (located on Chr4) served as control. Fold differences were calculated relative to strain M2. *NRG1* expression by Tri7 strains was ∼1.5-fold higher than that by euploid strains. *NRG1* expression by M2-Nrg1OE was∼1.8-fold higher than that of M2. **B.** Hyphal formation by strains was assessed on RPMI1640 with 2% glucose medium. Cells at exponential stage growth were spotted onto agar plates and photographed following incubation at 30°C for 5 days. Strain M2 formed normal hyphae whereas hyphal formation by M2-Nrg1OE and M1 was attenuated. **C.** Biofilm formation was assessed using a crystal violet assay. Data in panels A and C present results from 3 independent experiments; bars represent mean ± SEM. *P*-values were calculated using ANOVA with Dunnett’s multiple comparison tests, * *p*-values <0.05, ** *p* _≤_0.01, *** *p*_≤_0.001, **** *p* _≤_0.0001.

**Supplemental Figure 5.**
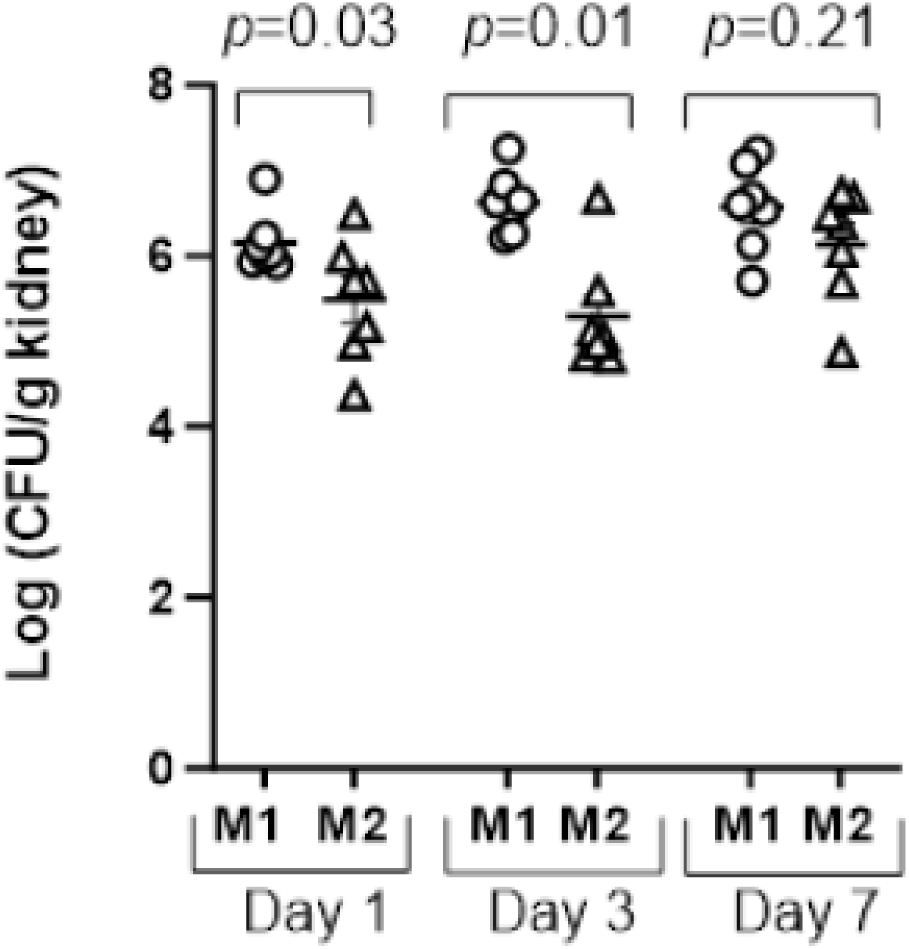
Tissue burdens of strains M1 and M2 within mouse kidneys during disseminated candidiasis. Mice were infected intravenously via lateral tail vein with either bar-coded M1 or M2 (1×10^6^ CFU). Kidneys were obtained for *C. albicans* burden enumeration. Data were log-transformed, and presented as mean ± SEM from 8 mice per strain for each timepoint. Burdens of strain M1 were significantly greater than those of strain M2 at d1 and d3 (*p*-values determined using Mann-Whitney for each timepoint).

**Supplemental Figure 6.**
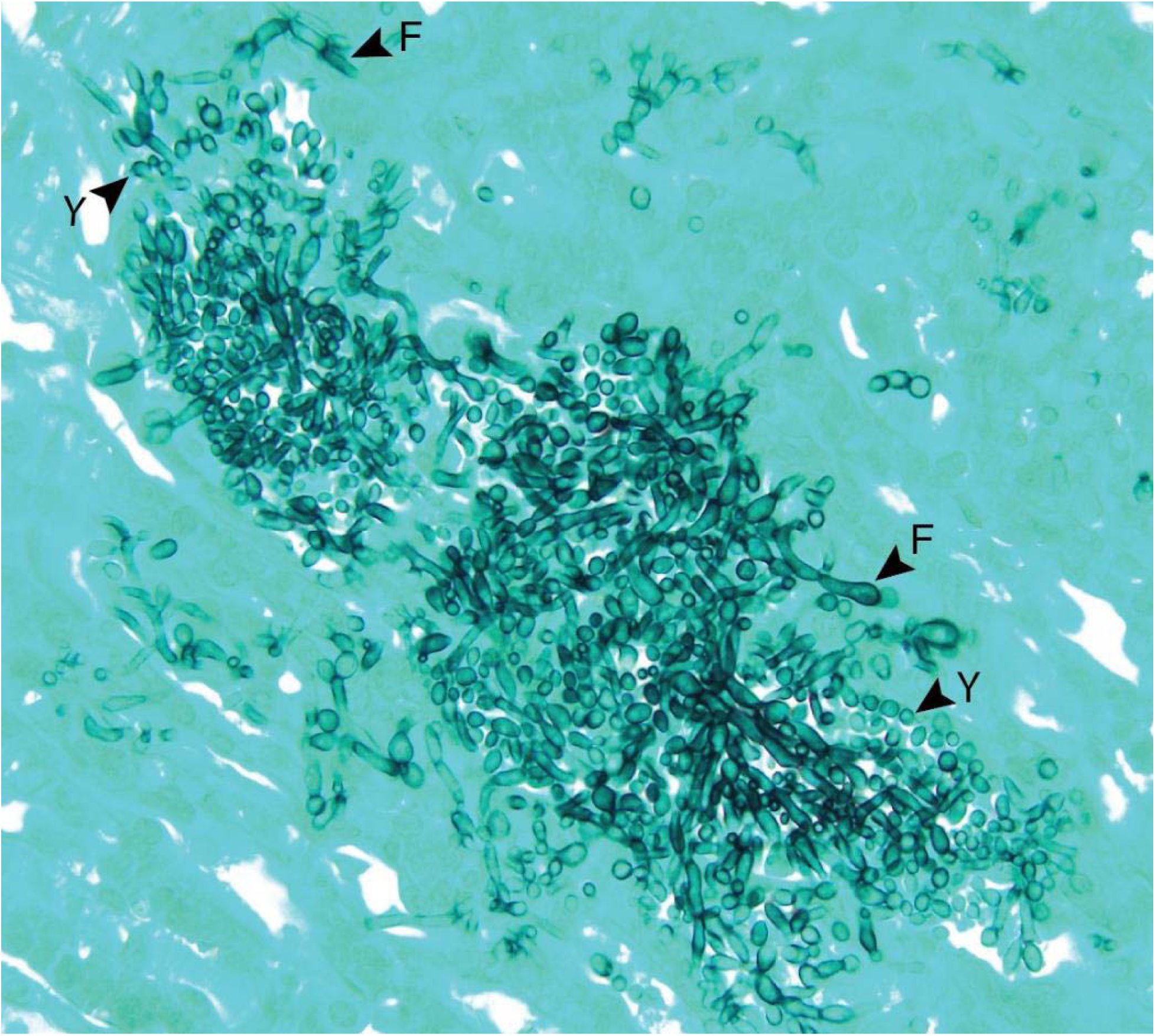
Morphology of strain M1 within mouse kidneys. Kidney sections from mice infected intravenously with barcoded strains M1 or M2 alone were stained with Grocott Methenamine Silver (GMS). Strain M1 is shown growing in filamentous morphologies (denoted by arrowheads labeled with “F”) interspersed with multiple yeasts (arrowhead labeled with “Y”) (image at 400x) within kidneys on d3. Filamentous morphologies included both hyphae and pseudohyphae. M1 morphologies on d1 were similar to those on d3, and they did not differ on either day from M2 morphologies (not shown).

**Supplemental Figure 7.**
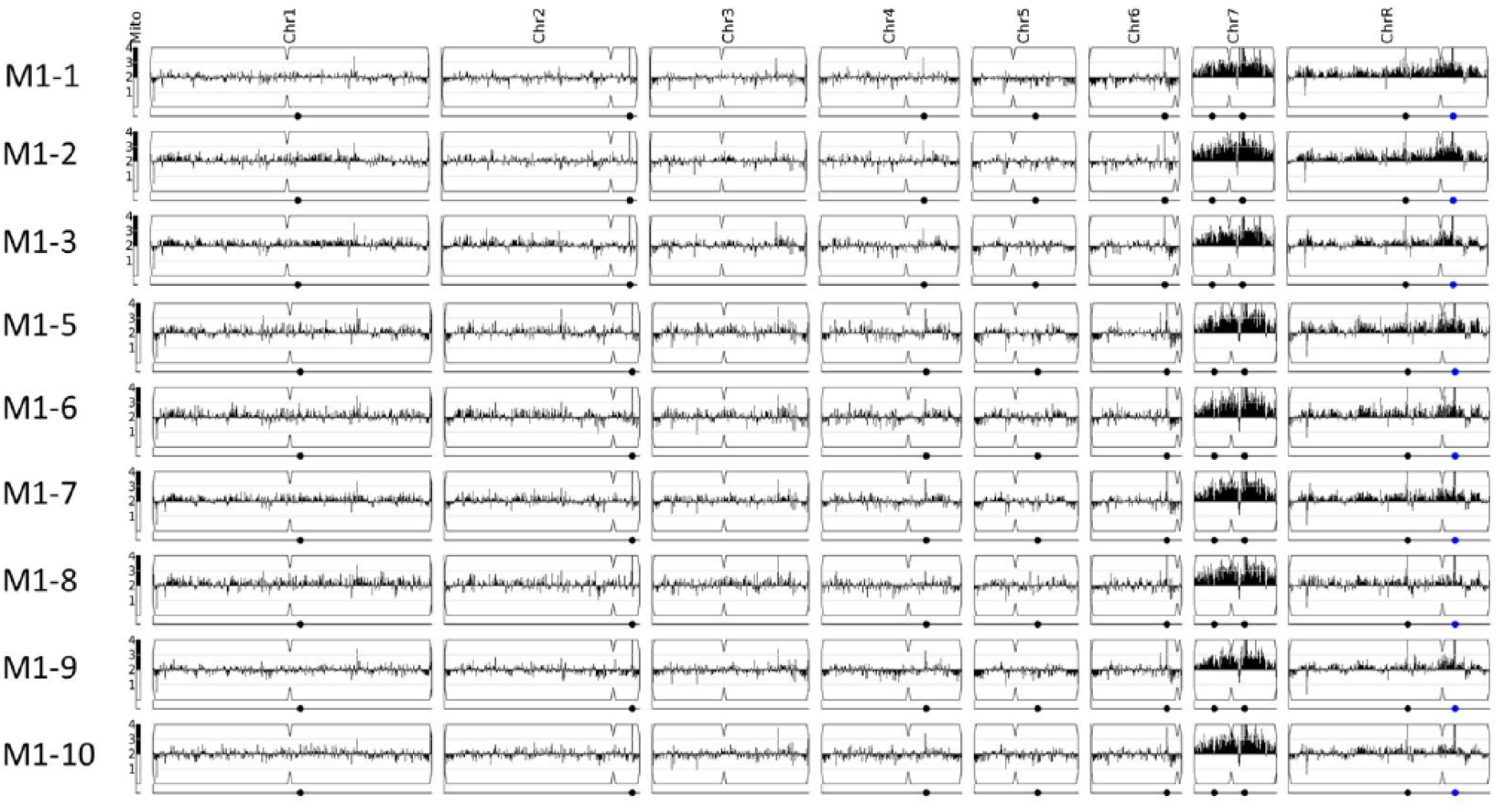
Copy number variation of 10 strains isolated from a blood culture bottle spiked with M1 index strain.

**Supplemental Table 1.**
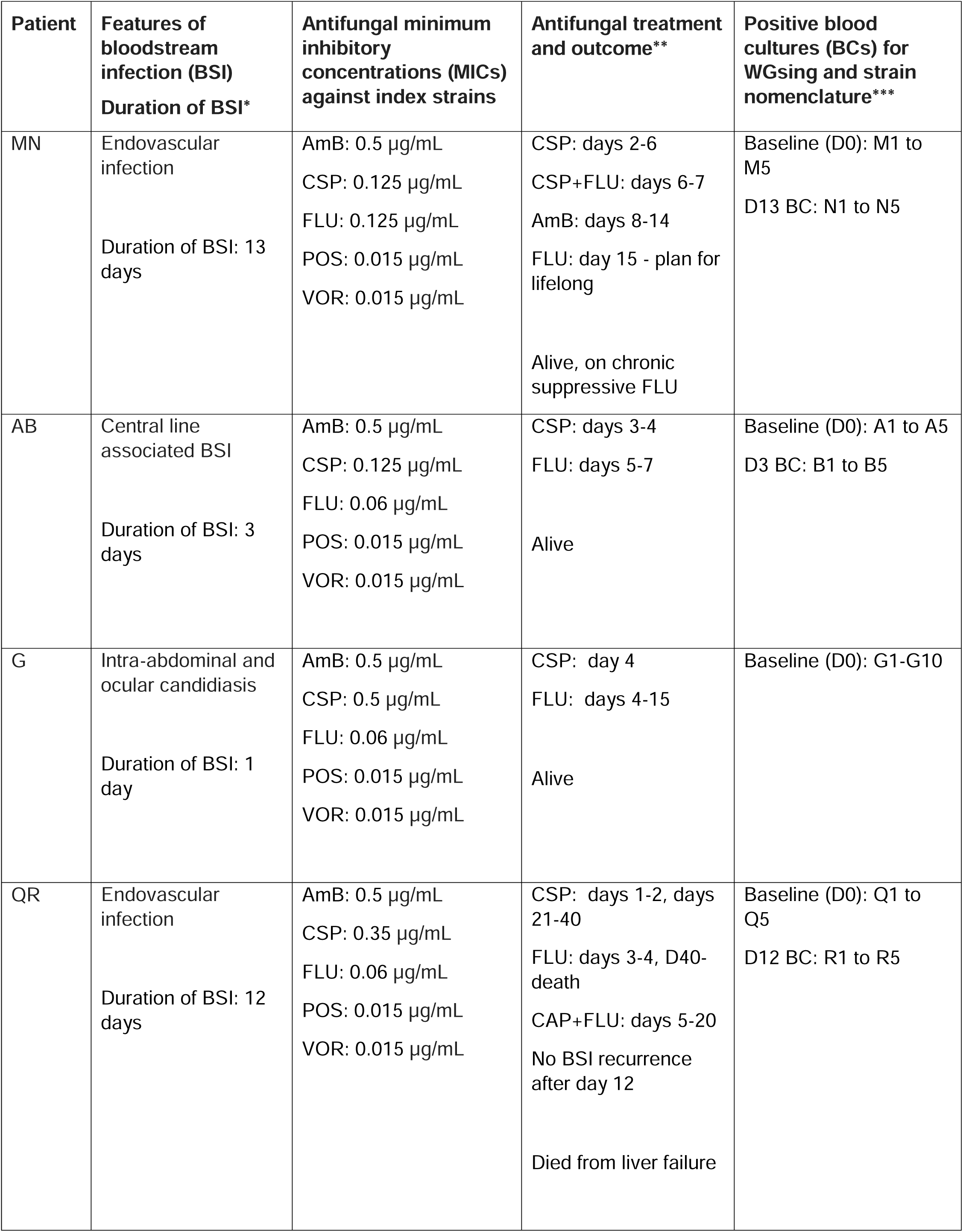
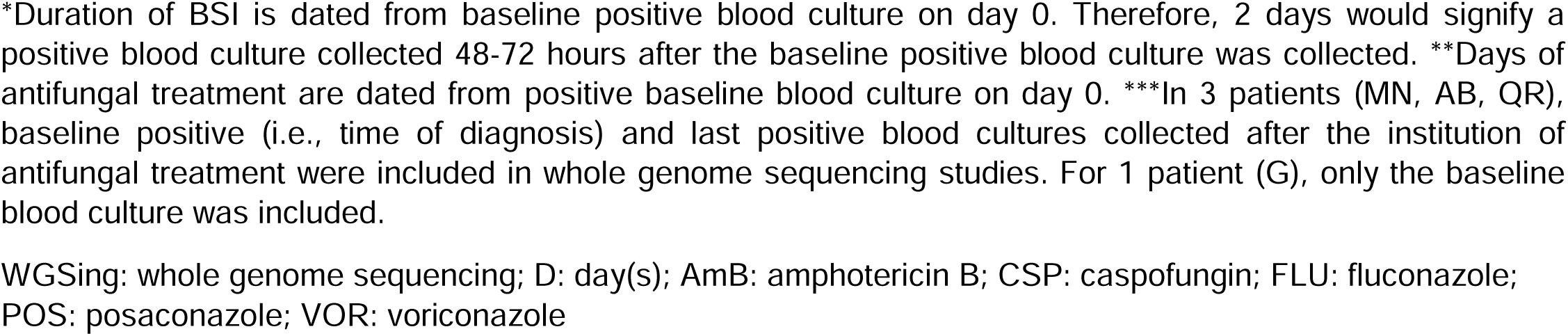
Clinical characteristics, treatment and outcomes of patients with *Candida albicans* bloodstream infections.

| Patient | Features of bloodstream infection (BSI)<br>Duration of BSI* | Antifungal minimum inhibitory concentrations (MICs) against index strains | Antifungal treatment and outcome** | Positive blood cultures (BCs) for WGsing and strain nomenclature*** |
| --- | --- | --- | --- | --- |
| MN | Endovascular infection<br>Duration of BSI: 13 days | AmB: 0.5 µg/mL<br>CSP: 0.125 µg/mL<br>FLU: 0.125 µg/mL<br>POS: 0.015 µg/mL<br>VOR: 0.015 µg/mL | CSP: days 2-6<br>CSP+FLU: days 6-7<br>AmB: days 8-14<br>FLU: day 15 - plan for lifelong<br>Alive, on chronic suppressive FLU | Baseline (D0): M1 to M5<br>D13 BC: N1 to N5 |
| AB | Central line associated BSI<br>Duration of BSI: 3 days | AmB: 0.5 µg/mL<br>CSP: 0.125 µg/mL<br>FLU: 0.06 µg/mL<br>POS: 0.015 µg/mL<br>VOR: 0.015 µg/mL | CSP: days 3-4<br>FLU: days 5-7<br>Alive | Baseline (D0): A1 to A5<br>D3 BC: B1 to B5 |
| G | Intra-abdominal and ocular candidiasis<br>Duration of BSI: 1 day | AmB: 0.5 µg/mL<br>CSP: 0.5 µg/mL<br>FLU: 0.06 µg/mL<br>POS: 0.015 µg/mL<br>VOR: 0.015 µg/mL | CSP: day 4<br>FLU: days 4-15<br>Alive | Baseline (D0): G1-G10 |
| QR | Endovascular infection<br>Duration of BSI: 12 days | AmB: 0.5 µg/mL<br>CSP: 0.35 µg/mL<br>FLU: 0.06 µg/mL<br>POS: 0.015 µg/mL<br>VOR: 0.015 µg/mL | CSP: days 1-2, days 21-40<br>FLU: days 3-4, D40-death<br>CAP+FLU: days 5-20<br>No BSI recurrence after day 12<br>Died from liver failure | Baseline (D0): Q1 to Q5<br>D12 BC: R1 to R5 |

\*Duration of BSI is dated from baseline positive blood culture on day 0. Therefore, 2 days would signify a positive blood culture collected 48-72 hours after the baseline positive blood culture was collected. \*\*Days of antifungal treatment are dated from positive baseline blood culture on day 0. \*\*\*In 3 patients (MN, AB, QR), baseline positive (i.e., time of diagnosis) and last positive blood cultures collected after the institution of antifungal treatment were included in whole genome sequencing studies. For 1 patient (G), only the baseline blood culture was included.
WGSing: whole genome sequencing; D: day(s); AmB: amphotericin B; CSP: caspofungin; FLU: fluconazole; POS: posaconazole; VOR: voriconazole

**Supplemental Table 2.**
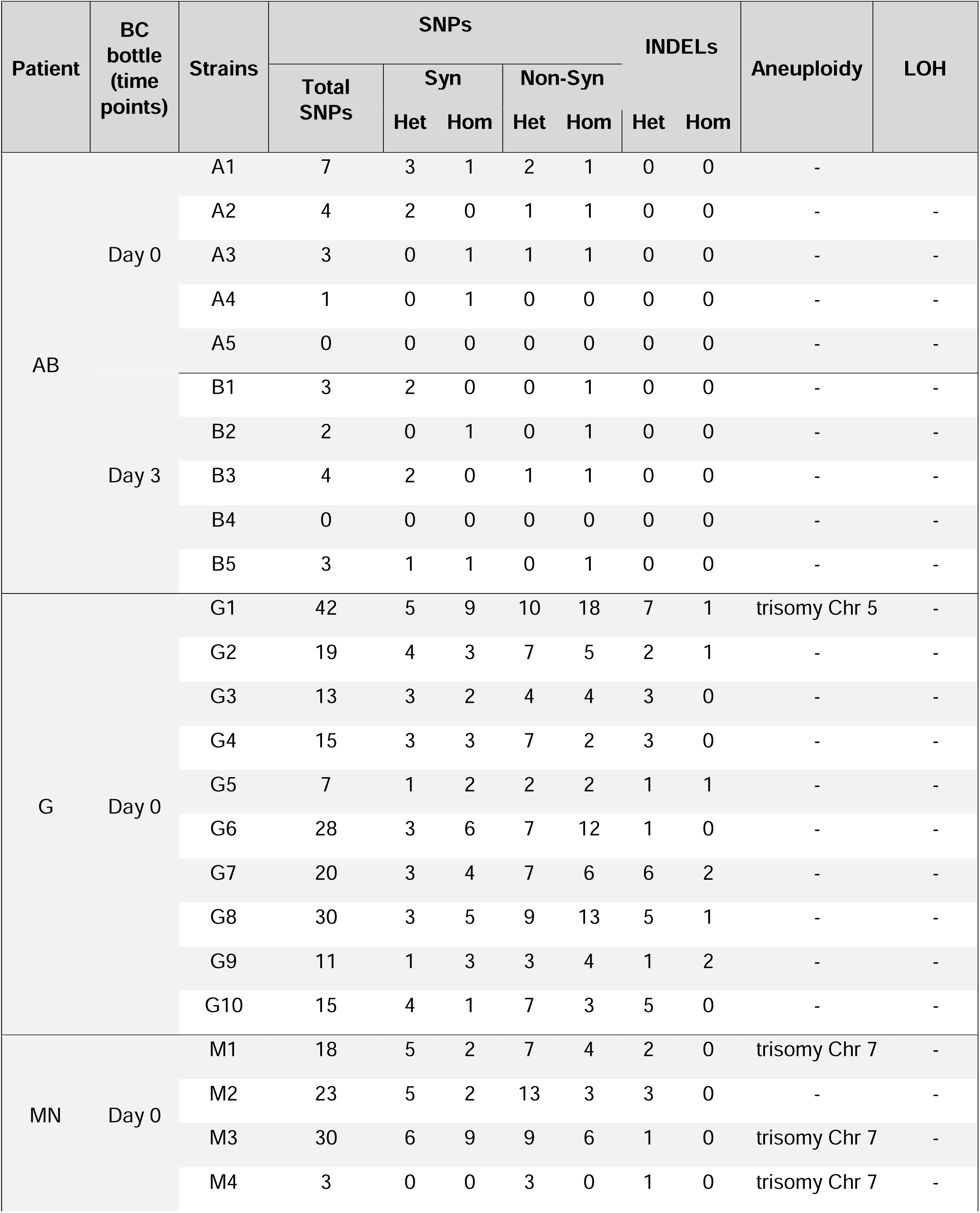
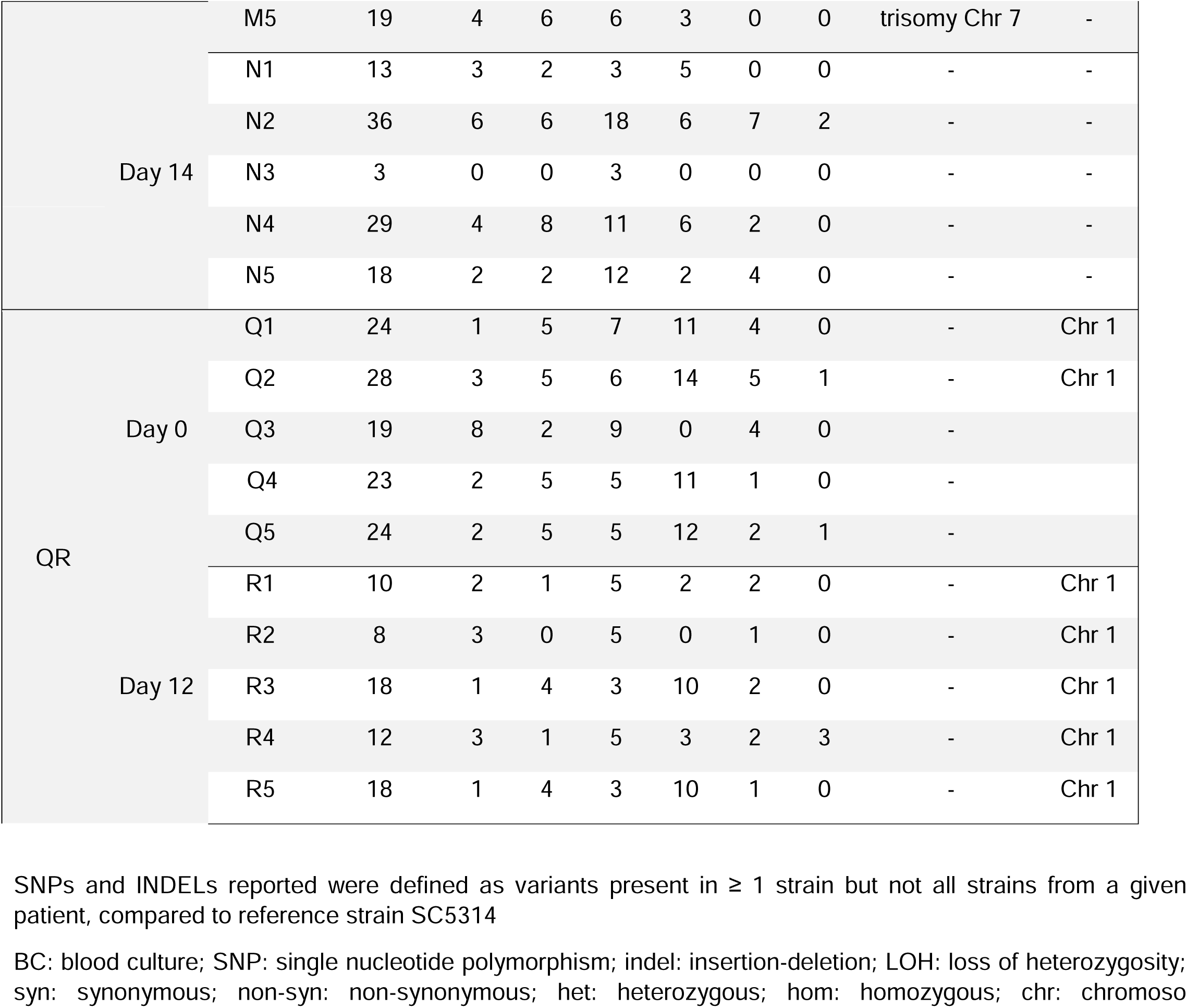
Within-patient genetic variations among *Candida albicans* strains from blood cultures.

**Supplemental Table 3.**
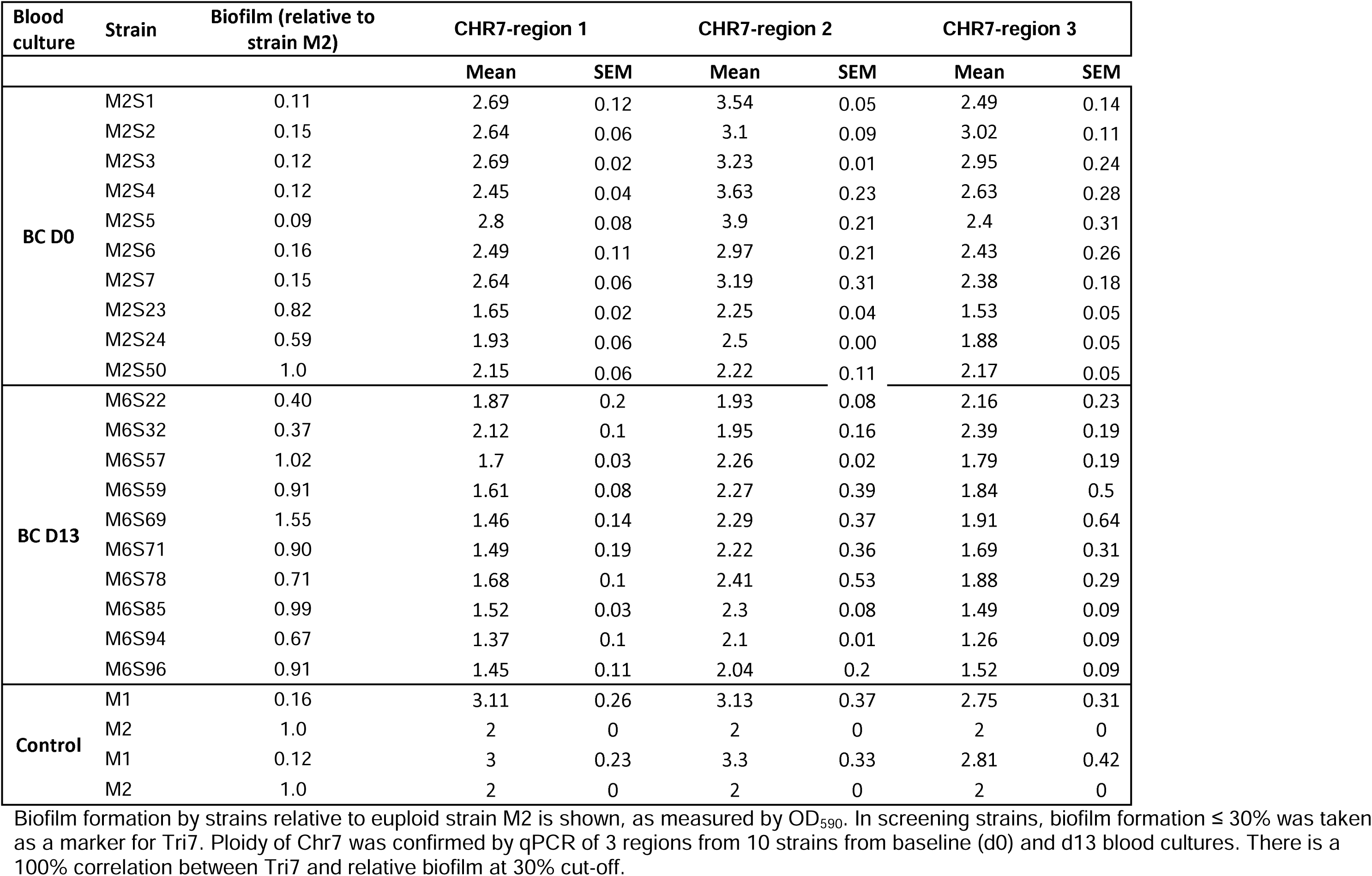
Validation of Tri7 among *C. albicans* strains recovered from blood cultures of patient MN.

| Blood culture | Strain | Biofilm (relative to strain M2) | CHR7-region 1 |  | CHR7-region 2 |  | CHR7-region 3 |  |
| --- | --- | --- | --- | --- | --- | --- | --- | --- |
|  |  |  | Mean | SEM | Mean | SEM | Mean | SEM |
| BC D0 | M2S1 | 0.11 | 2.69 | 0.12 | 3.54 | 0.05 | 2.49 | 0.14 |
|  | M2S2 | 0.15 | 2.64 | 0.06 | 3.1 | 0.09 | 3.02 | 0.11 |
|  | M2S3 | 0.12 | 2.69 | 0.02 | 3.23 | 0.01 | 2.95 | 0.24 |
|  | M2S4 | 0.12 | 2.45 | 0.04 | 3.63 | 0.23 | 2.63 | 0.28 |
|  | M2S5 | 0.09 | 2.8 | 0.08 | 3.9 | 0.21 | 2.4 | 0.31 |
|  | M2S6 | 0.16 | 2.49 | 0.11 | 2.97 | 0.21 | 2.43 | 0.26 |
|  | M2S7 | 0.15 | 2.64 | 0.06 | 3.19 | 0.31 | 2.38 | 0.18 |
|  | M2S23 | 0.82 | 1.65 | 0.02 | 2.25 | 0.04 | 1.53 | 0.05 |
|  | M2S24 | 0.59 | 1.93 | 0.06 | 2.5 | 0.00 | 1.88 | 0.05 |
|  | M2S50 | 1.0 | 2.15 | 0.06 | 2.22 | 0.11 | 2.17 | 0.05 |
| BC D13 | M6S22 | 0.40 | 1.87 | 0.2 | 1.93 | 0.08 | 2.16 | 0.23 |
|  | M6S32 | 0.37 | 2.12 | 0.1 | 1.95 | 0.16 | 2.39 | 0.19 |
|  | M6S57 | 1.02 | 1.7 | 0.03 | 2.26 | 0.02 | 1.79 | 0.19 |
|  | M6S59 | 0.91 | 1.61 | 0.08 | 2.27 | 0.39 | 1.84 | 0.5 |
|  | M6S69 | 1.55 | 1.46 | 0.14 | 2.29 | 0.37 | 1.91 | 0.64 |
|  | M6S71 | 0.90 | 1.49 | 0.19 | 2.22 | 0.36 | 1.69 | 0.31 |
|  | M6S78 | 0.71 | 1.68 | 0.1 | 2.41 | 0.53 | 1.88 | 0.29 |
|  | M6S85 | 0.99 | 1.52 | 0.03 | 2.3 | 0.08 | 1.49 | 0.09 |
|  | M6S94 | 0.67 | 1.37 | 0.1 | 2.1 | 0.01 | 1.26 | 0.09 |
|  | M6S96 | 0.91 | 1.45 | 0.11 | 2.04 | 0.2 | 1.52 | 0.09 |
| Control | M1 | 0.16 | 3.11 | 0.26 | 3.13 | 0.37 | 2.75 | 0.31 |
|  | M2 | 1.0 | 2 | 0 | 2 | 0 | 2 | 0 |
|  | M1 | 0.12 | 3 | 0.23 | 3.3 | 0.33 | 2.81 | 0.42 |
|  | M2 | 1.0 | 2 | 0 | 2 | 0 | 2 | 0 |

Biofilm formation by strains relative to euploid strain M2 is shown, as measured by OD<sub>590</sub>. In screening strains, biofilm formation ≤ 30% was taken as a marker for Tri7. Ploidy of Chr7 was confirmed by qPCR of 3 regions from 10 strains from baseline (d0) and d13 blood cultures. There is a 100% correlation between Tri7 and relative biofilm at 30% cut-off.

**Supplemental Table 4.** Strains and plasmids used in this study.

| Strains | Source | Strain description | Reference |
| --- | --- | --- | --- |
| Patient AB |  |  |  |
| A1 | Initial blood culture, d0 | Index strain, recovered by micro lab | This study |
| A2-A5 | Initial blood culture, d0 | Studied strains, recovered by PI lab |  |
| B6 | Subsequent blood culture, d12 | Index strain, recovered by micro lab |  |
| B7-10 | Subsequent blood culture, d12 | Studied strains, recovered by PI lab |  |
| Patient G |  |  |  |
| G1 | Initial blood culture, d0 | Index strain, recovered by micro lab | This study |
| G2-G10 | Initial blood culture, d0 | Studied strains, recovered by PI lab |  |
| Patient MN |  |  |  |
| M1 | Initial blood culture, d0 | Index strain, recovered by micro lab | This study |
| M2-M5 | Initial blood culture, d0 | Studied strains, recovered by PI lab |  |
| N6 | Subsequent blood culture, d13 | Index strain, recovered by micro lab |  |
| N7-10 | Subsequent blood culture, d13 | Studied strains, recovered by PI lab |  |
| Patient QR |  |  |  |
| Q1 | Initial blood culture, d0 | Index strain, recovered by micro lab | This study |
| Q2-Q5 | Initial blood culture, d0 | Studied strains, recovered by PI lab |  |
| R6 | Subsequent blood culture, d11 | Index strain, recovered by micro lab |  |
| R6-R10 | Subsequent blood culture, d11 | Studied strains, recovered by PI lab |  |
| Other strains |  |  |  |
| Barcoded-M1 | From M1 strain | PI lab | This study |
| Barcoded-M2 | From M2 strain | PI lab |  |
| NRG1- overexpressed in M2 | From M2 strain | PI lab |  |
| Plasmids |  |  |  |
| Plasmids RB793 | For barcoding of M1 | Addgene #199121 |  |
| Plasmid RB794 | For barcoding of M2 | Addgene #199122 |  |
| pCJN542 | provided by Mitchell AP | NRG1 over-expression |  |

**Supplemental Table 5.** Reagents and material used in this study.

|  |  |
| --- | --- |
| **Medium** |  |
| Sabouraud dextrose agar | Fisher Scientific, DF0109-07-07-3 |
| Yeast peptone dextrose (YPD) | Yeast Extract: BD Difco, 210929<br>Peptone: BD Difco, 11677<br>Dextrose: Fisher Scientific, D16-500 |
| Spider medium | Nutrient broth: Fisher Scientific, DF0003-17-8<br>Mannitol: Fisher Scientific, AC125345000<br>Potassium Phosphate Dibasic: Fisher Scientific, 7758-11-4<br>Agar, Fisher Scientific, DF0140-01-0 |
| RPMI 1640 medium | Sigma Aldrich, R1383 |
| **Chemicals** |  |
| Crystal violet | Fisher Scientific, C581-100 |
| DTT | Sigma Aldrich, 11583786001 |
| EDTA | Sigma Aldrich, E8008 |
| Glucose | Fisher Scientific, 50-99-7 |
| Phosphate buffer saline | Fisher Scientific, R582350010F |
| Sorbitol | Fisher Scientific, BP439-500 |
| **Reagents/enzymes** |  |
| DNaseI | New England Biolabs, M0303L |
| Lyticase | Fisher Scientific, NC1644419 |
| NgoMIV | New England Biolabs, R0564S |
| **Antimicrobial agents** |  |
| Ampicillin | Fisher Scientific, BP1760-25 |
| Chloramphenicol | Fisher Scientific, AC227921000 |
| Kanamycin | Fisher Scientific, AC450810500 |
| Nourseothricin | GoldBio, N-500-500 |
| Streptomycin | MP Biomedicals, 219454180 |
| Anidulafungin | MedChemExpress, HY13553 |
| Fluconazole | Fisher Scientific 86386-73-4 |
| Isavuconazole | BOC Sciences, 241479-67-4 |
| Micafungin | MedChemExpress, HY-16321 |
| Posaconazole | Fisher Scientific (TCI America products), P2477 |
| Voriconazole | Fisher Scientific (TCI America products) VO116 |
| **Kits** |  |
| PCR clean-up system | Promega, A9281 |
| Phusion High-Fidelity PCR mix with HF buffer | New England Biolabs, M0531S |
| Qiagen's DNeasy Blood and Tissue Kit | Qiagen (Hilden, Germany), 69504 |
| qRT-PCR mix | Fisher Scientific, FERK0223 |
| RiboPure RNA purification kit for yeast | Invitrogen, AM1924 |
| SYBR Green qPCR Master Mix | Fisher Scientific, FERK0223 |
| Verso cDNA synthesis kit | Fisher Scientific, AB1453A |
| Yeast DNA extraction kit | ThermoFisher, 788707 |
| Zymobiomics DNA Miniprep kit | ZymoResearch, D4300 |
| **Others** |  |
| FBS (Fetal Bovine Serum) | HyClone, SH30071.02 |
| Sterile culture bottle BD BACTEC Plus Aerobic medium | Fisher Scientific, BD442023 |

**Supplemental Table 6.**
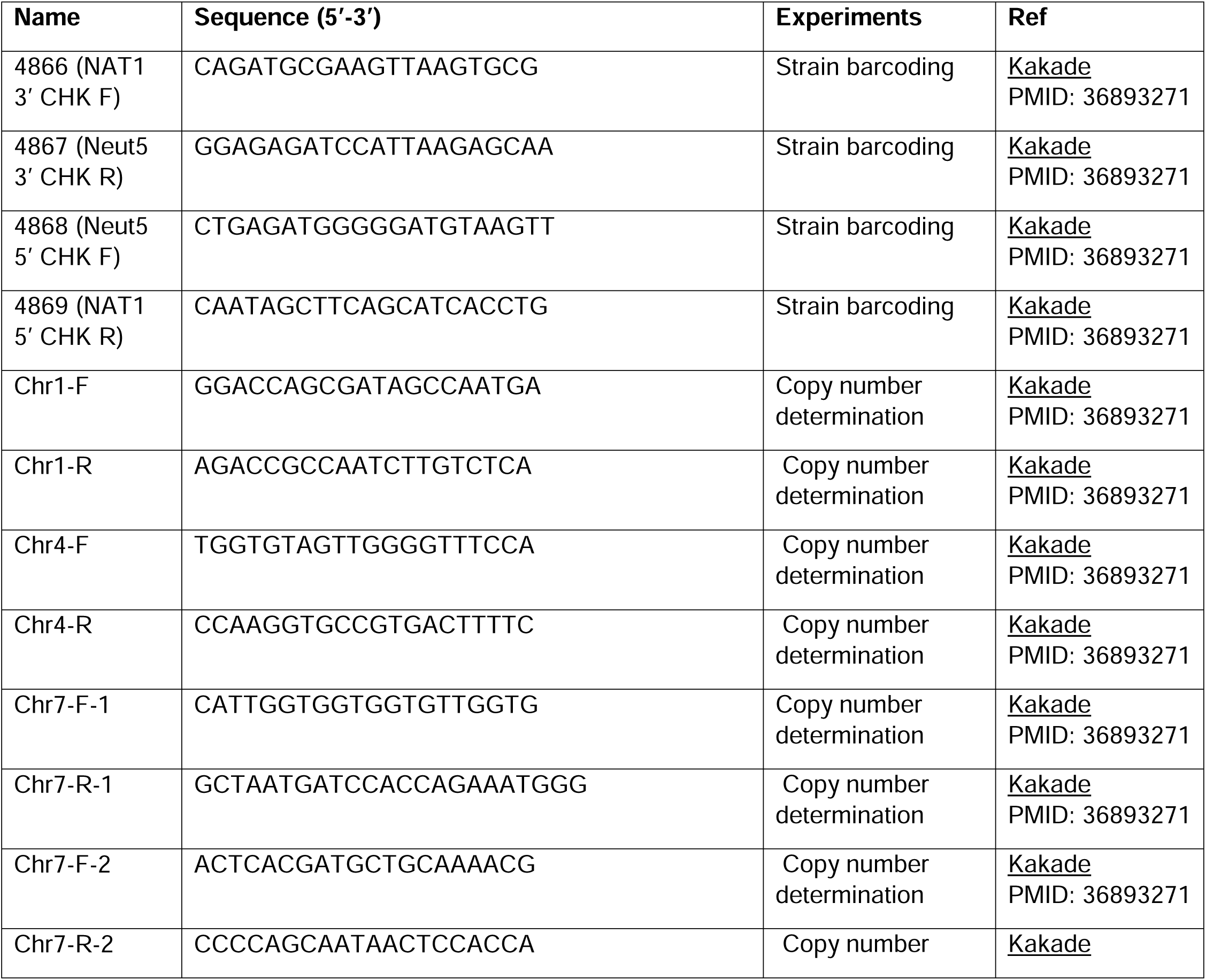
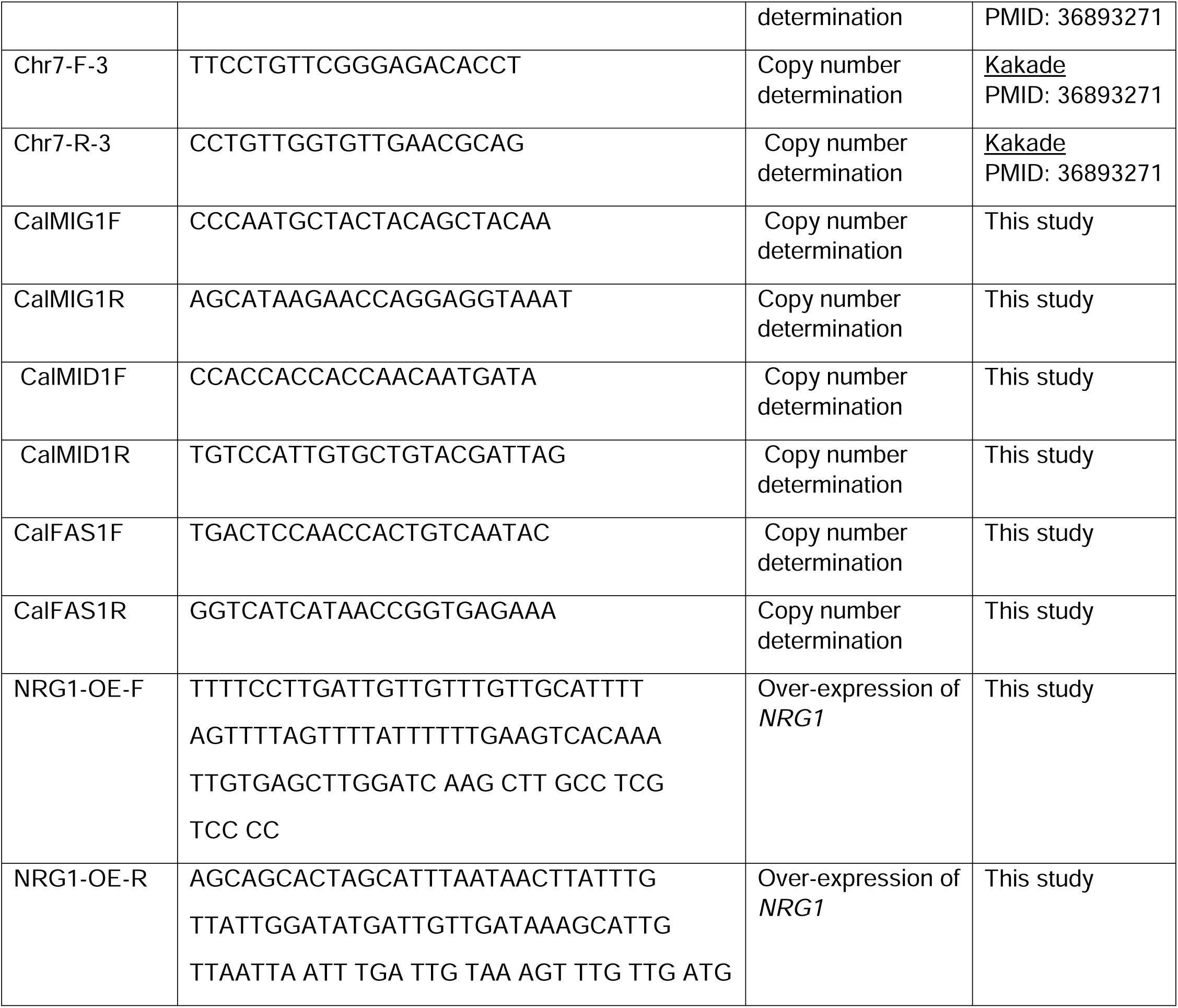
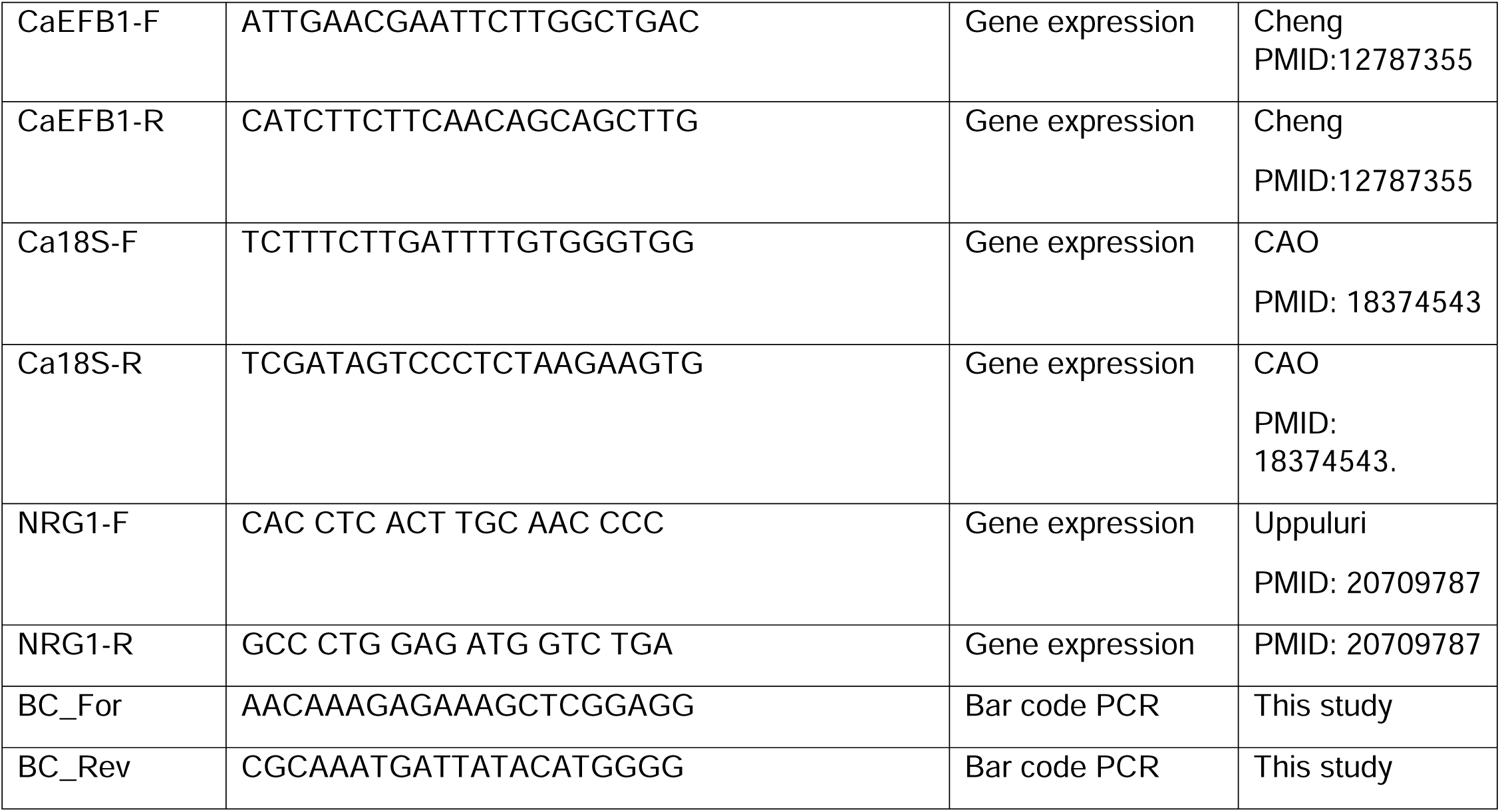
Primers used in this study.

## Bioinformatic analyses

DNA was extracted using Qiagen’s DNeasy Blood and Tissue Kit (Hilden, Germany).(Badrane, Cheng et al. 2023) DNA libraries were prepared with the Nextera XT (Illumina, San Diego) protocol and sequenced to ≥ 50X coverage with 2×150 bp paired end reads. Raw read zipped files were analyzed through a custom-built NextFlow pipeline, which integrates Burrows-Wheeler Aligner for reads mapping to the *C. albicans* SC5314 genome downloaded from Candida Genome Database (http://www.candidagenome.org/). The Genome Analysis Toolkit (GATK, v4.5.0.0) (McKenna, Hanna et al. 2010) was used for deduplicating and sorting of mapped reads, variant calling and filtering after a base quality score recalibration. SnpEff v4.5 was employed for variant annotation.(Cingolani, Platts et al. 2012) Low-confidence variants were filtered using GATK *VariantFiltration* tool. Variants were included if they fulfilled ≥ 1 of 4 stringent statistical criteria (i.e., variants were filtered if they failed all 4 criteria). The compound filtering expression was: Quality by Depth (QD) < 2; Fisher Strand (FS) > 60; mean square Mapping Quality (MQ) < 40. Additional filtering was applied through customized filtering criteria: minimum genotype quality (GQ) of 50; minimum of 80% support to the alternate allele in allele depth (AD), and minimum depth (DP) of 10. After filtration, the select variants tool extracted SNPs for final VCF files. VCF variant files from all strains were merged using BCFTOOLS.(Danecek, Bonfield et al. 2021) Whole genome multiple sequence alignment (MSA) was extracted from the multi-sample VCF file using VCF2MSA python script (https://github.com/tkchafin/vcf2msa.py). Fasta aligned sequences for each contig were extracted from the merged VCF file using a python script (https://github.com/Bahler-Lab/alignment-from-vcf). All aligned contigs were then merged and converted to a phylip format using a perl script (https://github.com/nylander/catfasta2phyml). A phylogenomic tree was built using RaxML(Kozlov, Darriba et al. 2019) and visualized in iTol v6.(Letunic and Bork 2021)

## *In vitro* phenotypic assays

### In vitro growth

*C. albicans* strains were grown in liquid YPD medium overnight at 30°C and diluted in YPD or RPMI1640 with 2% dextrose to OD_600_ (Optical Density, 600 nm) of 0.1. Diluted cultures were added to 96-well plates in triplicate with 200 μL per well, and growth was monitored at 30°C for 24h in a microplate reader (Molecular Device). Growth curves were analyzed using GraphPad Prism 9. Experiments were performed with 3 biological replicates.

### *In vitro* filamentation

Strains were grown overnight in liquid YPD at 30°C and then cultured in YPD, YPD containing 10% Fetal Bovine Serum (FBS), RPMI1640 with 2% glucose or Spider media at 37 °C for 4 h with shaking (200 rpm). Cell suspensions were removed hourly and imaged using microscopy. For filamentation on solid medium, overnight culture grown in YPD at 30°C was washed and adjusted to 10^7^ cells/mL in normal saline; 5 μl were spotted on agar plate and incubated at 30°C.

### Antifungal susceptibility testing and tolerance assays

The sensitivity to cell wall stress-inducing agents were assessed by spotting 5 µL of serial ten-fold dilutions of each strain grown overnight in YPD onto SDA plates impregnated with the indicated chemical agent. The plates were incubated at 30 and 40°C until colonies appeared. Antifungal susceptibility testing was performed using the broth dilution technique according to the Clinical and Laboratory Standards Institute (CLSI) standardized method.((CLSI)) Concentrations ranged from 0.25-256□μg/mL for fluconazole (Fisher Scientific 86386-73-4), and 0.015-16□μg/mL for echinocandins. At 24 hours, the wells were read visually with a viewing mirror. MIC was set at the lowest drug concentration at which there was a ≥ 50% decrease in growth, relative to growth in no drug containing wells. The plates were also were read using a spectrophotometer reader (SPECTROstar, BMG labtech)) at OD600nm for determination of tolerance. Supra-MIC growth (SMG) was calculated as the average growth per well above the MIC divided by the level of growth without drug.(Rosenberg, Ene et al. 2018, Berman and Krysan 2020) Tolerance was also quantified using disk diffusion.(Rosenberg, Ene et al. 2018, Berman and Krysan 2020) For disk diffusion, overnight growth *C. albicans* cells (2□×□10^5^ CFU) were spread onto casitone plates (9□g/l Bacto casitone, 5□g/l yeast extract, 15□g/l Bacto agar, 11.5□g/l sodium citrate dehydrate and 2% glucose) with a cotton swab and allowed to air dry for ∼30 min. A filter disk containing micafungin (5ug) was placed in the center of the plate and allowed to grow at 35°C for 48 hours. The radius corresponding to the point where growth was inhibited by 20%, 50% or 80% relative to the maximum radius, was measured. Tolerance levels were determined from the degree of growth within the region of growth inhibition detected visually on the plates and calculated using *diskImageR* and imageJ (fraction of growth, FoG_20_, FoG_50_, FoG_80_, respectively).

## Biofilm formation

Biofilms were formed in flat-bottomed 96-well microplates. For each strain, a cell suspension in normal saline was adjusted to McFarland 0.5 and 1:10 diluted in RPMI1640 medium supplemented with 2% (w/v) glucose. Plate wells were inoculated with 200 μL of standardized *C.albicans* suspension in triplicate and incubated at 37 °C for 90 minutes to allow cell adhesion. A negative control was prepared by inoculating 200 μL RPMI1640 medium. After the adhesion phase, non-adherent cells were removed by thoroughly washing the wells with 0.15 M sterile phosphate-buffered saline (PBS, pH 7.2). Each well was then filled with 200 μL of fresh RPMI1640 medium, and plates were incubated at 37°C for 24h to allow biofilm formation. To assess biofilm formation, culture broth was gently aspirated, and each well was washed twice with PBS and incubated with 100□μL methanol for 15 min at room temperature. Plates were left to dry completely in a chemical safety cabinet, followed by staining with 100□μL 0.1% crystal violet for 5 min and three washes with dH_2_O. Crystal violet was dissolved from the stained biomass by adding 100□μL of 33% acetic acid and by plate shaking for 1□min at 800□rpm. Supernatants of dissolved crystal violet were transferred into fresh wells of a 96-well plate, and absorbance at 590□nm was recorded using a micro plate reader (Molecular Device).

